# The phosphatidylglycerol phosphate synthase PgsA utilizes a trifurcated amphipathic cavity for catalysis at the membrane-cytosol interface

**DOI:** 10.1101/2021.06.27.450103

**Authors:** Bowei Yang, Hebang Yao, Dianfan Li, Zhenfeng Liu

## Abstract

Phosphatidylglycerol is a crucial phospholipid found ubiquitously in biological membranes of prokaryotic and eukaryotic cells. The phosphatidylglycerol phosphate (PGP) synthase (PgsA), a membrane-embedded enzyme, catalyzes the primary reaction of phosphatidylglycerol biosynthesis. Mutations in *pgsA* frequently correlate with daptomycin resistance in *Staphylococcus aureus* and other prevalent infectious pathogens. Here we report the structures of *S. aureus* PgsA (*Sa*PgsA) captured at two distinct states of the catalytic process, with lipid substrate (cytidine diphosphate-diacylglycerol, CDP-DAG) or product (PGP) bound to the active site within a trifurcated amphipathic cavity. The hydrophilic head groups of CDP-DAG and PGP occupy two different pockets in the cavity, inducing local conformational changes. An elongated membrane-exposed surface groove accommodates the fatty acyl chains of CDP-DAG/PGP and opens a lateral portal for lipid entry/release. Remarkably, the daptomycin resistance-related mutations mostly cluster around the active site, causing reduction of enzymatic activity. Our results provide detailed mechanistic insights into the dynamic catalytic process of PgsA and structural frameworks beneficial for development of antimicrobial agents targeting PgsA from pathogenic bacteria.

## Introduction

Phospholipids and membrane proteins are the major components of biological membrane [1]. In prokaryotic and eukaryotic cells, phospholipids fulfill indispensable roles by contributing to formation of biological membranes and participating in various fundamental biochemical processes [2, 3]. They are amphipathic molecules with hydrophobic fatty acyl chains and hydrophilic head groups.

Glycerophospholipids are the dominant type of phospholipids with multitudinous structures and functions [4]. Among various phospholipids with different head groups, phosphatidylglycerol (PG) is a pivotal anionic phospholipid present in bacteria, plants and animals [5]. PG and its derivatives (aminoacyl PG and cardiolipin/CL) are the most abundant phospholipids in *Staphylococcus aureus*, a Gram-positive bacterium [6]. Meanwhile, PG and CL are the major anionic phospholipids in *Escherichia coli* (a Gram-negative bacterium) membrane besides phosphatidylethanolamine/PE, the most abundant zwitterionic phospholipid in *E. coli* [7]. In plants, PG is the only major phospholipid in the photosynthetic membranes of chloroplasts and has important functions for the electron transport process during photosynthesis, chloroplast development and chilling tolerance [8]. In mammals, PG is crucial for lung function as it is one of the major phospholipids found in lung surfactants and modulates the surface activity of lung, and it may also regulate innate immunity and inhibit infections caused by respiratory viruses [9-11]. Besides, PG is also involved in activation of RNA synthesis and nuclear protein kinase C as well as inhibition of platelet activating factor and phosphatidylcholine (PC) transfer in mammalian cells [5].

As the PG precursor, phosphatidylglycerol phosphate (PGP) is synthesized through the cytidine diphosphate-diacylglycerol (CDP-DAG)−dependent pathway, a fundamental phospholipid biosynthesis process presents in both prokaryotic and eukaryotic cells [12, 13]. The bacterial PGP synthase PgsA is a membrane-embedded enzyme that converts CDP-DAG and glycerol 3-phosphate (G3P) into PGP and cytidine monophosphate (CMP) at the interface between membrane and cytosol [14] (Figure 1A). To further generate PG, PGP is dephosphorylated by phosphatases to remove the terminal phosphate group [15]. Subsequently, PG can be utilized by the cardiolipin synthase as the substrate to produce CL [7]. Alternatively, it can be used by the aminoacyl phosphatidylglycerol synthases to produce aminoacyl PG (aaPG), an important class of PG derivatives crucial for bacteria to adapt to environmental changes and resist cationic antimicrobial peptides or antibiotics [6, 16].

**Figure 1.**
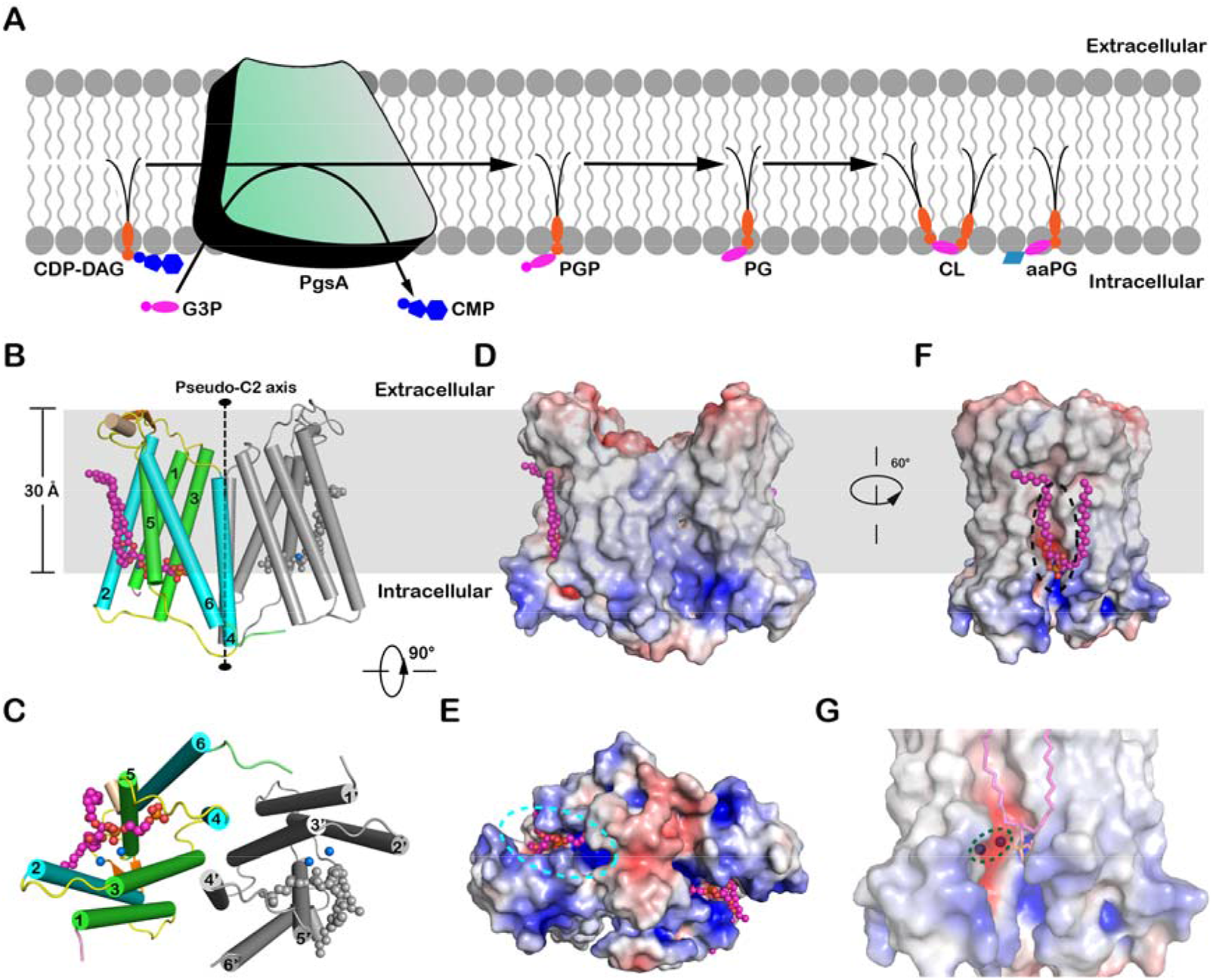
Biological function and overall structure of *Sa*PgsA. (**A**) A cartoon diagram showing the functional role of PgsA in biosynthesis of PG and its derivatives in the membrane. (**B** and **C**) The overall structure of *Sa*PgsA dimer shown in cartoon model viewed along membrane plane (**B**) or membrane normal from cytoplasmic side (**C**). Color codes: green, TM1, TM3 and TM5; cyan, TM2, TM4 and TM6; violet, the N-terminal region; lime, the C-terminal region; yellow, loop regions between adjacent TMs; orange, the β-hairpin in the TM1-TM2 loop region; wheat, the short α-helix between TM5 and TM6. The lipid molecules (PGP) are shown as magenta or gray sphere models, whereas the Zinc ions are presented as marine blue spheres. For clarity, one protomer is shown in colored mode, whereas the adjacent protomer is in light grey. (**D**-**F**) The electrostatic potential surface representation of *Sa*PgsA. The views are along the membrane plane from two different angles (**D**, **F**) or along membrane normal from intracellular side (**E**). Color codes: blue, electropositive; white, neutral; red, electronegative. The grey box in the background of **B**, **D** and **F** indicates the approximate position of the hydrophobic region of membrane according to the result of prediction through the Positioning of Proteins in Membrane (PPM) server. The cyan dashed elliptical ring in **E** indicates the position of active-site cavity within the *Sa*PgsA monomer, while the black dashed elliptical ring in **F** labels the approximate location of a lateral portal connecting the active site with lipid bilayer. (**G**) A close-up view of the active site center with a bimetal binding site. The green dashed elliptical ring indicates the location of two metal ions in the active site of *Sa*PgsA.

Dysfunctional mutation of *pgsA* leads to drastic reduction of PG, CL and PG derivatives in *E. coli* membrane which are essential for cell growth and viability [17]. Moreover, *pgsA* is frequently associated with daptomycin resistance in pathogenic bacteria posing threats to human health [18-20]. Daptomycin is a cyclic lipopeptide antibiotic used for treating serious infections caused by methicillin-resistant *S. aureus* (MRSA), vancomycin-resistant *Enterococci* (VRE) and other Gram-positive pathogens [21]. Accumulating evidences show that mutations in *pgsA* correlate with daptomycin resistance in several Gram-positive bacteria such as *S. aureus*, *Bacillus subtilis*, *Corynebacterium striatum*, *Staphylococcus capitis* and *Streptococcus oralis* [18-20, 22-25]. Furthermore, PgsA from *S. aureus* (*Sa*PgsA) has recently been identified as a potential antibacterial target to eradicate methicillin-resistant *S. aureus* (MRSA) persisters [26].

Cyanobacteria with dysfunctional PgsA exhibit impaired assembly, oligomerization and function of both photosystem I (PSI) and photosystem II (PSII) [27]. Besides, PGP synthase gene expression is regulated by factors that affect mitochondrial development, making the enzyme an excellent indicator for mitochondrial membrane biogenesis [28]. A previous study found that the activity of mitochondrial PGP synthase was increased by 21% and 98% in 15- and 22-month-old rats respectively, in comparison to the 2-month-old animals suffering from spontaneously hypertensive heart failure [29]. Moreover, a recent study proposed that mitochondrial cardiolipin synthases might evolve from prokaryotic PgsA through the neofunctionalization of the bacterial ancestor [30].

Despite that PgsA has important biological functions and is closely associated with antibiotic resistance of pathogenic bacteria, little is known about its structure, catalytic mechanism and functional link between PgsA mutations and daptomycin resistance. To shed light into the catalytic mechanism of PgsA and assist the future development of antibacterial agents, we have solved the crystal structures of *Sa*PgsA with PGP and CDP-DAG bound to the active site at resolutions of 2.5 Å and 3.0 Å, respectively. Moreover, the daptomycin resistance-related mutations are mapped onto the structure and enzymatic activity assays have been carried out to characterize the functional effect of the mutations. Structure-based mutagenesis and enzymatic assay results allow us to propose a working model for PGP biosynthesis. Meanwhile, our high-resolution structures of *Sa*PgsA may also serve as a framework for rational design of antimicrobial drugs.

## Results

### Structure determination and overall structural features of *Sa*PgsA

Purified *Sa*PgsA was reconstituted into lipid cubic phase (LCP), a lipid-based matrix providing membrane-like environment for crystallization of membrane proteins [31]. The initial phases of *Sa*PgsA−PGP complex structure were solved through the *ab initio* phasing and chain tracing method (Supplementary Fig. 1A and B) [32]. *Sa*PgsA molecules were packed in two-dimensional lattices, which further stacked orderly in the third dimension through interlayer contacts to form the type I membrane-protein crystal (Supplementary Fig. 1C). The structures of *Sa*PgsA were refined to 2.5 Å (PGP−bound state) and 3.0 Å (CDP-DAG−bound state) resolution, respectively (Supplementary Table 1). For clarity, the higher resolution *Sa*PgsA−PGP complex structure is used for overall fold analysis.

As shown in Figure 1B and 1C, *Sa*PgsA forms a homodimer with two protomers related by a central pseudo two-fold symmetry axis and each protein monomer forms a compact fold with six transmembrane α-helices (TMs 1-6). The *Sa*PgsA dimer is not only observed in the crystal grown in LCP [33], but is also present in detergent solution (Supplementary Fig. 1D). The dimerization interface of *Sa*PgsA is stabilized mostly through the extensive hydrophobic interactions between TM3 and TM4’ (the prime symbol indicates TMs from the adjacent protomer) and between TM4 and TM4’ in the membrane-embedded region (Figure 1B and 1C).

While PgsA belongs to the CDP-alcohol phosphatidyltransferase (CDP-OH_P_transf) superfamily (http://pfam.xfam.org/family/CDP-OH_P_transf), the overall structure of *Sa*PgsA differs from the other members of CDP-OH_P_transf superfamily with known structures as it shares fairly low sequence identity (∼20%) with them (Supplementary Fig. 2A and C). CDP-OH_P_transf enzymes are all integral membrane proteins catalyzing the scission of a phosphoric anhydride bond to release CMP from a CDP-alcohol and concomitant formation of a new phosphodiester bond in the presence of divalent cation [3]. They contain a strictly conserved motif (D_1_xxD_2_G1xxAR…G_2_xxxD_3_xxxD_4_) that is crucial for catalysis (Supplementary Fig. 2A and B) [34-38]. As proposed previously [36], CDP-OH_P_transf superfamily can be further classified into three families (A, B and C) basing on membrane topology and evolutionary relationship (Supplementary Fig. 3). Most of prokaryotic PgsAs belong to family A which has not been structurally characterized. The mechanism of product formation, binding and release steps in the catalytic process mediated by CDP-OH_P_transf enzyme remains unclear.

Analysis through the Positioning of Proteins in Membrane (PPM) server [39] indicates that most of the *Sa*PgsA is embedded in the membrane (Figure 1B and 1D). The amino-terminal region of *Sa*PgsA is fairly short with only two residues. In comparison, the structures of phosphatidylglycerol phosphate synthase (PIPS) from *Renibacterium salmoninarum* and *Mycobacterium tuberculosis* possess much longer N-terminal regions forming a juxta-membrane helix [36, 37]. On the extracellular side, all six TMs of *Sa*PgsA are positioned underneath the membrane surface (Figure 1B), creating a concave surface which may induce local membrane deformation (Figure 1D). Two extracellular regions, namely the long TM1-2 loop and the TM5-6 loop with a short four-residue α-helix, are positioned near the membrane surface, whereas a much shorter TM3-4 loop is located below the membrane surface (Figure 1B and 1C). On the intracellular side, all TMs of *Sa*PgsA including the shortest one (TM5) extend out of the membrane surface and protrude into cytoplasm (Figure 1B). For the structures of other CDP-OH_P_transf members, such as AF2299 and PIPS from *Mycobacterium kansasii* [35, 38], TM5s are all completely buried in the membrane instead (Supplementary Fig. 2C).

Curiously, a large open cavity is harbored in the membrane-embedded core of *Sa*PgsA (Figure 1E). This cavity, surrounded by TM2, TM3, TM4 and TM5, has an amphipathic surface (Figure 1D and 1F). While TM4 and TM5 are straight α-helices, TM2 and TM3 each contains a conserved kink vital for enzymatic activity [34]. The cavity has a hydrophilic portal connected with the cytoplasm and a lateral hydrophobic portal opening toward the membrane (Figure 1E and 1F). Such a feature differs from the one in the di-*myo*-inositol-1,3’-phosphate-1’phosphate synthase from *Archaeoglobus fulgidus* (*Af*DIPPS) [34], as *Af*DIPPS contains two small pockets opening to the cytoplasm to acquire and accommodate two hydrophilic substrates.

The intracellular entrance of *Sa*PgsA is mainly shaped by the TM2-3 and TM4-5 loops located on the intracellular surface, while the lateral membrane-exposed portal is located at the groove between TM2 and TM5. The lateral portal may serve as the gateway for hydrophobic substrate to enter the cavity from the membrane or for the lipid product to return to the membrane. Similarly, the hydrophilic substrate (G3P) or product (CMP) might use the intracellular entrance to access or exit from the active site. In support of this hypothesis, the surface around the intracellular entrance is overall electropositive and may thus facilitate the entry of substrate and release of product via electrostatic attraction.

### An amphipathic cavity with an electropositive pocket for PGP binding

To understand the mechanism of product formation, binding and release process, we have solved the structure of *Sa*PgsA in complex with its product PGP (Supplementary Table 1, *Sa*PgsA/PGP). The 2*F*_o_-*F*_c_ and the omit electron density maps fit well with the structural model of PGP (Supplementary Fig. 4A). Consistently, mass spectrometry analysis on the lipid sample extracted from purified *Sa*PgsA protein revealed characteristic peaks of intact PGP and its fragments (Supplementary Fig. 4B). It is noteworthy that PGP was co-purified along with *Sa*PgsA protein and preserved during protein purification and crystallization process.

In the *Sa*PgsA−PGP complex structure, each protomer binds one endogenous PGP molecule in the active site (Figure 2A). The PGP binding site is embedded mostly in the hydrophobic region and harbored inside the large intracellular cavity. The hydrophilic head group of PGP is positioned nearby the bimetal catalytic center within the electropositive region of the cavity exposed to the cytoplasm (Figure 2A). The 3’-phosphoryl moiety of PGP is sandwiched between TM3 and TM5, and interacts with the evolutionarily conserved Lys83, Arg110, Arg118, Lys137 and Tyr181 (Figure 2A and Supplementary Fig. 2B). The positively charged Arg118 and hydroxyl-containing Tyr181 are close to the intracellular surface of membrane, and may attract the negatively charged substrate G3P into the active site. A characteristic feature of the PGP binding site lies in the strongly electropositive region made of three positively charged residues (Lys83, Arg110 and Lys137). Such arrangement is favorable for neutralizing the negative charges on the G3P head group of PGP. Consistently, K83A, R110A, R118A and K137A mutants exhibited activity lower than the wild type. In contrast, the Y181A mutation increased the activity, as the mutation may facilitate entry of G3P into the active site by opening the pathway wider (Figure 2A and 2B). Besides, the backbone 3-phosphoryl moiety of PGP is located between TM2 and TM5 (Figure 2A). Thr138 on TM5 forms a hydrogen bond with the 2’-hydroxyl of PGP, and the loss of this interaction may facilitate PGP release from the active center. Therefore, T138A exhibited slightly higher activity than the wild type (Figure 2B). On the other hand, the 1-acyl chain of PGP extends along the groove between TM2 and TM5, whereas the 2-acyl chain is positioned on the hydrophobic surface of TM5 and TM6 (Figure 2C). Since CMP is absent in the *Sa*PgsA−PGP complex structure, it may represent an intermediate state of the catalytic process when the product PGP has formed and after CMP has been released.

**Figure 2.**
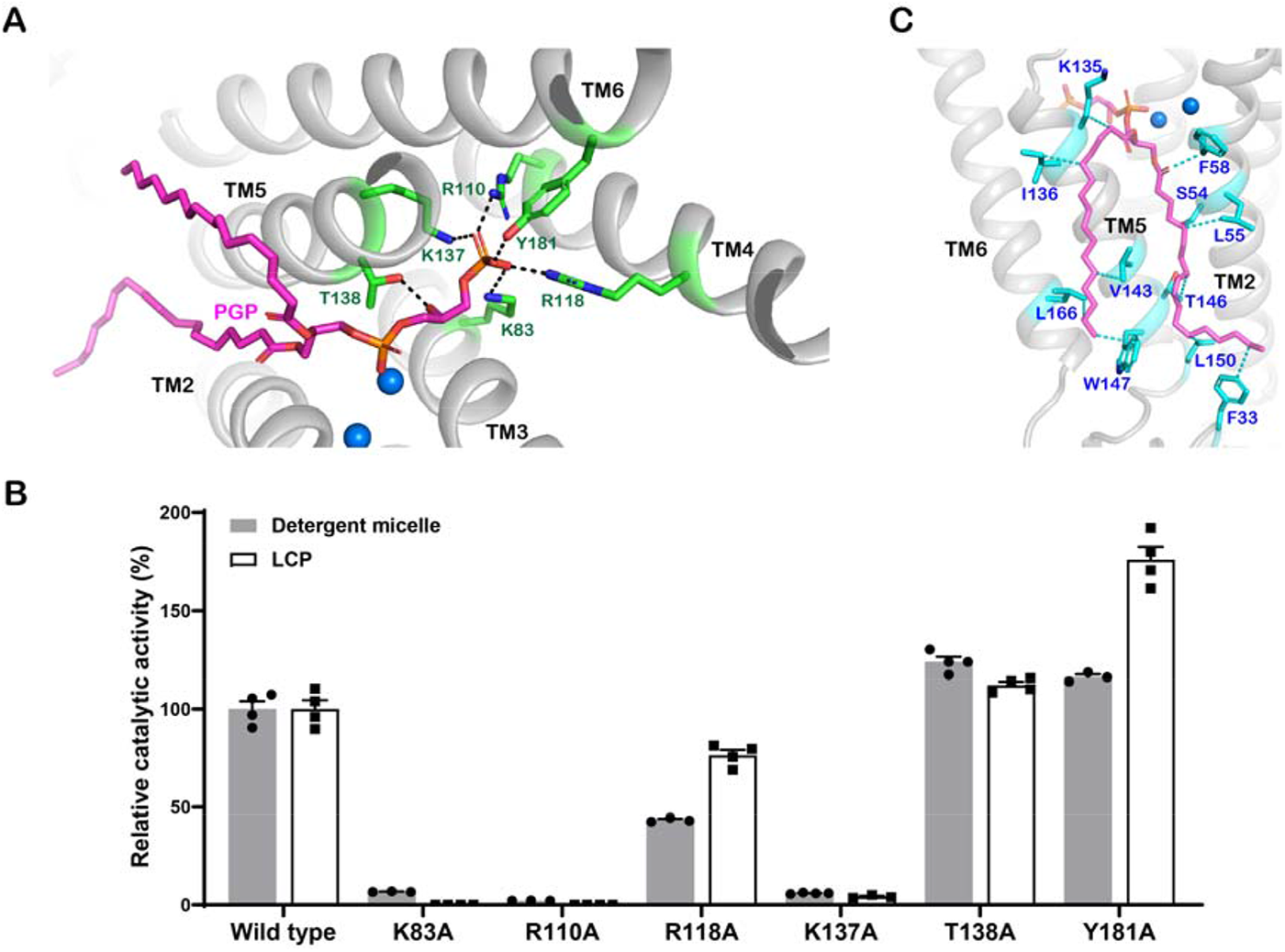
The PGP binding site in *Sa*PgsA. (**A**) Interactions between PGP and *Sa*PgsA through salt bridges and hydrogen bonds. Six evolutionarily conserved residues are coloured in green. Black dashed lines indicate the polar interactions between PGP head group and adjacent residues with bond lengths at 2.3-3.0 Å. The two blue spheres indicate two metal ions in the active. (**B**) The relative catalytic activity of six point mutants related to the PGP−binding site. The error bars indicate ± SEM with n = 3 or 4. The solid grey bars represent the activity data measured with WT and mutant enzymes solubilized in detergent, whereas the white bars represent the data measured with enzymes embedded in LCP. (**C**) Hydrophobic and van der Waals interactions between the acyl chains of PGP and *Sa*PgsA. Relevant residues are colored in cyan. The cyan dashed lines indicate the interactions between PGP acyl chains and adjacent residues at distances of 3.2-3.9 Å.

### The CDP-DAG-binding site partially overlaps with that of PGP

To unravel the substrate binding sites of *Sa*PgsA, we performed crystallization trials of the enzyme in the presence of both CDP-DAG and G3P at a molar ratio of 1:10:40 (*Sa*PgsA: CDP-DAG: G3P). Although the amount of G3P added into the crystallization mixture is in excess relative to the protein and CDP-DAG molecule, no recognizable G3P densities were observed in the active site. Nevertheless, the 2*F*_o_-*F*_c_ and omit electron density maps show clear densities for CDP-DAG (Supplementary Fig. 4C), and one CDP-DAG molecule is found in the active site of each monomer.

The CDP moiety of CDP-DAG interacts with Asn5 and Arg65 through hydrogen bonds and salt bridges, and the nucleotide ring wedges into a pocket delineated by TM1, TM2, TM3 and TM2-3 loop (Figure 3A). Among the eight highly conserved amino acid residues (D_57_xxD_60_G_61_xxA_64_R_65_…G_74_xxxD_78_xxxD_82_) in the active site of *Sa*PgsA, Gly61, Ala64, Arg65 and Gly74 outline the pocket for accommodating the nucleotide moiety of CDP-DAG. The two glycine residues may provide local structural flexibility for the release of CMP. A wide opening surrounded by electropositive surface extends from the binding site of the CDP-DAG nucleotide ring to the cytosol, potentially serving as the exit portal for CMP (Figure 3B, Channel 1). The acyl chains of CDP-DAG are located at the groove between TM2 and TM5, like those of PGP observed in the *Sa*PgsA−PGP complex structure (Figure 3C and 3D). The binding site may allow the enzyme to acquire CDP-DAG from membrane through lateral diffusion.

**Figure 3.**
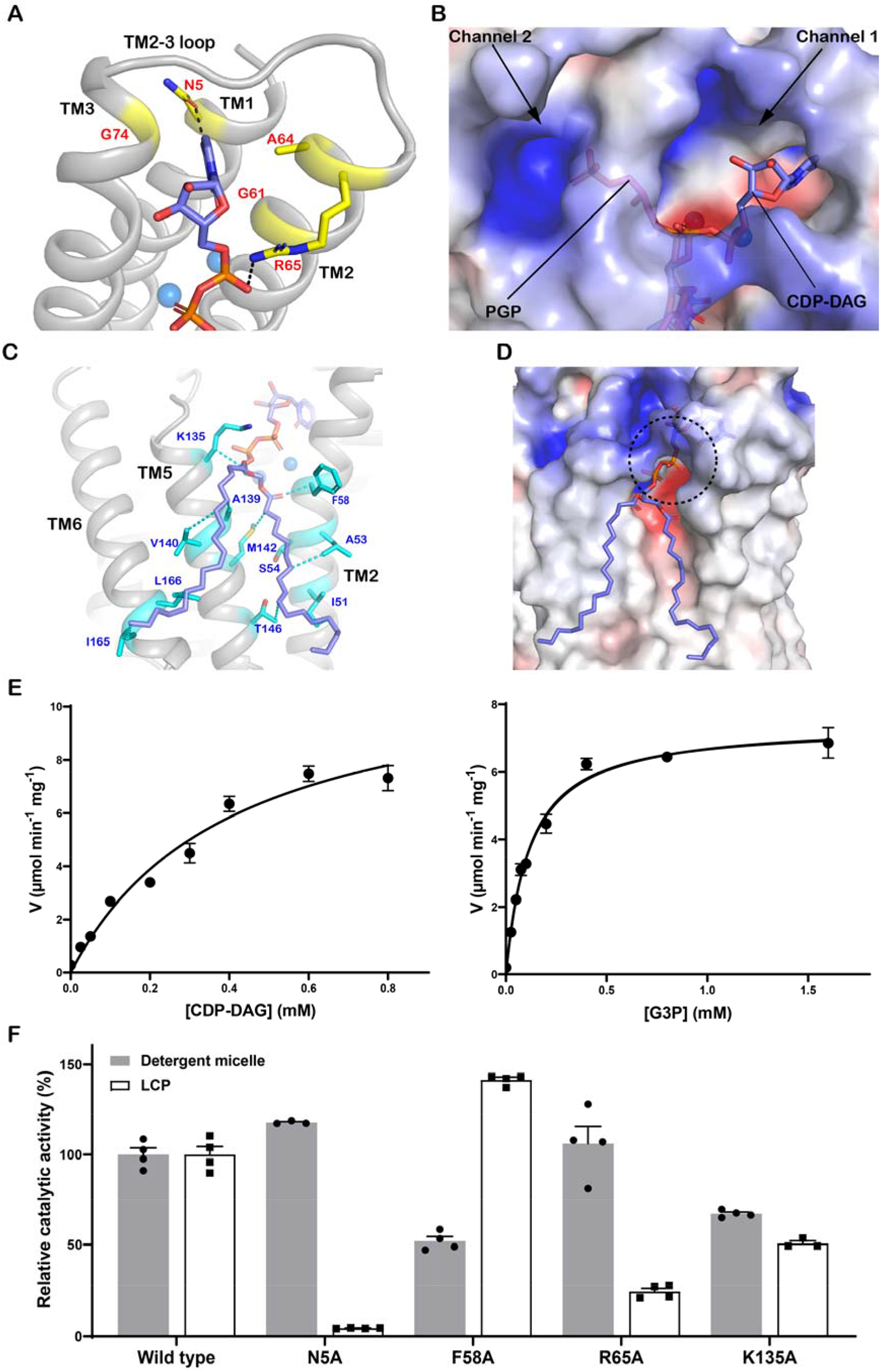
The binding site of CDP-DAG in *Sa*PgsA. (**A**) Interactions of the CDP moiety of CDP-DAG with *Sa*PgsA. The black dashed lines indicate hydrogen bonds and salt bridges at bond lengths of 2.4 Å or 3.1 Å. (**B**) Putative CMP release channel (Channel 1) and G3P entrance channel (Channel 2). The PGP molecule from the *Sa*PgsA−PGP complex structure is superposed onto the *Sa*PgsA−CDP-DAG complex structure to indicate the putative G3P location. (**C**) Interaction of the fatty acyl chains of CDP-DAG with adjacent amino acid residues in *Sa*PgsA. The relevant residues are highlighted in cyan. The cyan dashed lines indicate hydrophobic and van der Waals interactions at distance of 3.1-4.0 Å. (**D**) The putative entrance for CDP-DAG head group as indicated by the black dashed circle. (**E**) Substrate-dependent kinetic measurement of *Sa*PgsA activity in detergent micelle. The data points are plotted as mean value ± standard error (SEM as indicated by the error bars. n=3 or 4). For those data points with small errors, the error bars are buried within the symbols. (**F**) The relative catalytic activity of four alanine mutants related to the CDP-DAG−binding site. The error bars indicate ± SEM with n = 3 or 4. The solid grey bars represent the activity data measured with WT and mutant enzymes solubilized in detergent, whereas the white bars represent the data measured with enzymes embedded in LCP.

The [CDP-DAG] and [G3P]-dependent kinetic assay results indicate *Sa*PgsA has *K*m values of 0.40 mM for CDP-DAG and 0.12 mM for G3P (Figure 3E, measured in detergent micelle). The enzyme exhibits little or no substrate-induced cooperativity as the Hill coefficient was close to 1 (0.90 and 1.04 for CDP-DAG and G3P respectively), suggesting the two protomers likely perform catalytic function independently. To be consistent with the crystallization environment, we also reconstituted *Sa*PgsA into LCP and assessed its activity using a direct and continuous spectroscopic assay (Supplementary Fig. 5A) [40]. *Sa*PgsA was active in the LCP bilayer, supporting the view that *Sa*PgsA was properly folded in LCP and that the structures were of functional relevance (Supplementary Fig. 5). The [G3P]-dependent (*K*_m_ of 0.37 mM) kinetics in LCP closely resemble those measured in detergent (Supplementary Fig. 5C). Likewise, the PGP-binding site mutants showed similar relative activities (*vs* wild type) in detergents or LCP (Figure 2B). Curiously, the [CDP-DAG]-dependent kinetic profile differs from the one measured in detergent (Supplementary Fig. 5C). While the exact reason for this difference is unclear, the results are not unexpected considering the dramatic differences between the two types of local environments around the enzyme. Putatively, the activity difference may reflect slower substrate-diffusion in the viscous LCP or that the enzyme acquires and converts the lipid substrate at different rates in different environments.

For the four residues (Asn5, Phe58, Arg65 and Lys135) interacting directly with CDP-DAG, the corresponding alanine mutants behave very differently in detergents and LCP (Figure 3F). In detergents, F58A and K135A exhibited reduced activity, while N5A and R65A showed activity close to the wild type. Considering that CDP-DAG can diffuse easily in the detergent-based environments, N5A and R65A may have no significant effect on the process of CDP-DAG access to the active-center. By contrast, N5A and R65A showed much lower activity than the wild type in LCP. In LCP, diffusion of CDP-DAG is restricted in two dimensions, presumably at a slower rate than in detergents. The enzyme may rely on Asn5 and Arg65 to attract and position CDP-DAG head group in the active site. In addition, the charge interaction between Lys135 and the α-phosphoryl moiety of CDP-DAG was also important, probably by steering the substrate to a proper position for catalysis. Phe58 is proximal to the fatty acyl chains of CDP-DAG and at the lateral portal for CDP-DAG entry (Figure 3C). The F58A mutation may change the local structure, potentially influencing the access of CDP-DAG. As Phe58 is exposed on a hydrophobic surface, the activity of F58A mutant appears strongly dependent on the local environments, showing higher relative activity (*vs* the wild type) in LCP than in detergent (Figure 3F).

### A bimetal binding site is located at the active-site center

Divalent cations are essential for the activity of CDP-OH_P_transf enzymes [34, 41, 42]. Previous studies indicated PgsA activity is highly dependent on Mg^2+^ [40, 41, 43]. As shown in Figure 4A and Supplementary Fig. 5B, the activity of *Sa*PgsA is highest in the condition with Mg^2+^, whereas it becomes less active in the presence of Zn^2+^, Cd^2+^, Co^2+^ or Mn^2+^ and inactive with Ba^2+^, Ca^2+^ or EDTA. Besides, *Sa*PgsA activity is inhibited by increasing concentration of Zn^2+^ with the half maximal inhibitory concentration (IC50) at ∼1.8 μM (Figure 4B). Notably, the *Sa*PgsA crystals were grown in high concentrations (0.2 or 0.3 M) of Zn^2+^. The structures of *Sa*PgsA contain two metal ions as suggested by the 2*F*_o_-*F*_c_ electron density map, they were identified as zinc ions according to the anomalous difference Fourier density map (Supplementary Fig. 6). The two Zn^2+^-binding sites in *Sa*PgsA structures are validated by the metal-ligand geometry and valence analysis result (Supplementary Fig. 7) [44]. A previous theoretical study indicated that Zn^2+^ can substitute Mg^2+^ directly in its octahedral binding site or readjust into a tetrahedral geometry during the exchange [45]. Moreover, Zn^2+^ and Mg^2+^ share similar electronegativity and ionic radius (Zn^2+^, 0.88 Å; Mg^2+^, 0.86 Å). Therefore, the binding mode of Mg^2+^ should closely resemble those of Zn^2+^ ions observed in the *Sa*PgsA structures.

**Figure 4.**
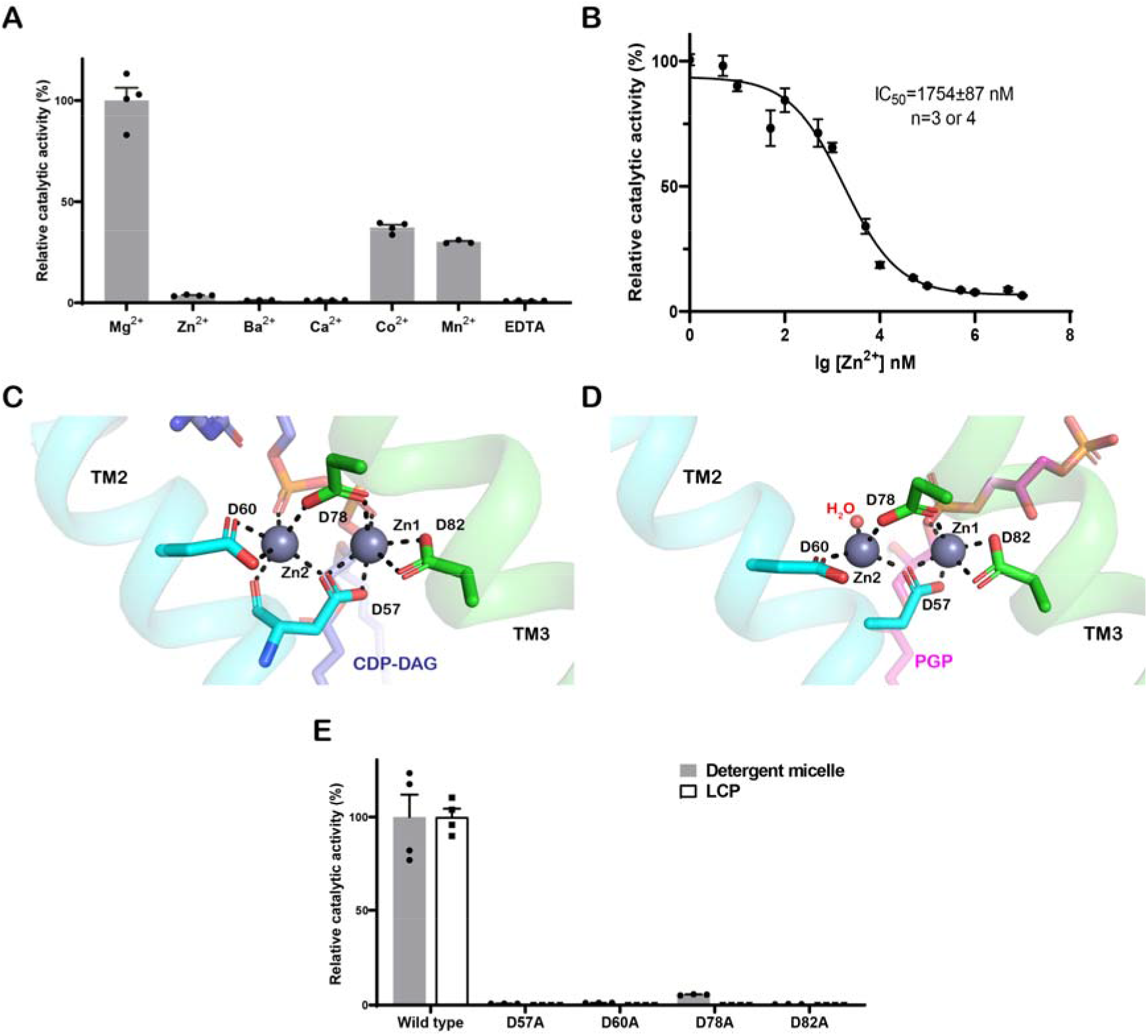
The bimetal center in *Sa*PgsA. (**A**) The dependence of PGP synthase activity of *Sa*PgsA on different divalent metal ions. (**B**) Inhibitory effect of Zn^2+^ concentration on *Sa*PgsA enzymatic activity in detergent micelle. The error bars indicate ± SEM with n = 3 or 4. For those data points with small errors, the error bars are buried within the symbols. (**C** and **D**) Detailed view of the bimetal binding sites in the *Sa*PgsA−CDP-DAG complex structure (**C**) and the *Sa*PgsA−PGP complex structure (**D**). Asp57 and Asp60 on TM2 are depicted as cyan sticks, while Asp78 and Asp82 on TM3 are coloured in green. CDP-DAG is shown as light blue stick, whereas PGP is shown as magenta stick. Black dashed lines indicate the coordination bonds at lengths of 2.0-2.3 Å. (**E**) The relative catalytic activity of metal-binding site mutants compared to the wild type. In (**A**) and (**E**), the error bars indicate ± SEM with n = 3 or 4. The solid grey bars represent the activity data measured with WT and mutant enzymes solubilized in detergent, whereas the white bars represent the data measured with enzymes embedded in LCP.

The *Sa*PgsA structures contain two zinc ions (Zn1 and Zn2) in the active site center. Zn1 and Zn2 are located at the bottom surface of the membrane-embedded cavity, and close to the lateral portal (Figure 1G). They are coordinated by Asp57, Asp60, Asp78 and Asp82 from a conserved sequence motif (D_57_xxD_60_G_61_xxA_64_R_65_…G_74_xxxD_78_xxxD_82_) at bond lengths of 2.0-2.3 Å (Supplementary Fig. 6). Meanwhile, Zn1 and Zn2 are bridged by Asp57 with bidentate coordination constraining the inter-zinc distance to 3.5 Å or 3.7 Å. Such a bimetal center structure tends to increase the p*K*_a_ of surrounding residues, thereby creating an acidic anion microenvironment to facilitate bond formation [34]. In detail, Zn1 is coordinated by Asp57, Asp78, Asp82 and the β-phosphoryl moiety of CDP-DAG, while Zn2 is coordinated by Asp57, Asp60, Asp78 and the α-phosphoryl moiety of CDP-DAG (Figure 4C). Similarly, in the *Sa*PgsA−PGP complex structure, a water molecule and the backbone 3-phosphoryl moiety of PGP replace the α-phosphoryl and β-phosphoryl moieties of CDP-DAG respectively (Figure 4D). Evidently, Zn1 coordinates the β-phosphoryl moiety of CDP-DAG in a proper position for the nucleophilic attack launched by the 1-OH of G3P. On the other hand, Zn2 may be involved in the scission of the α-β phosphoric anhydride bond of CDP-DAG, an important step to secure the irreversibility of catalysis [30]. The four acidic residues (Asp57, Asp60, Asp78 and Asp82), well positioned for coordination of Zn^2+^ and Zn^2+^-mediated CDP-DAG binding, are indispensable for PgsA activity, as removal of the metal-coordinating carboxyl group of the acidic residues by alanine mutation abolished enzyme activity (Figure 4E).

### *Sa*PgsA exhibits distinct local conformations at the two different states

Strikingly, the structure of *Sa*PgsA−PGP complex differs from that of *Sa*PgsA−CDP-DAG complex in terms of local conformations. While the common phosphatidyl moieties align relatively well, the CMP head group of CDP-DAG and the G3P head group of PGP point to nearly opposite directions (Figure 5A). As a result, the side chain of Asn5 on TM1 in the CDP-DAG−bound state moves outward by ∼1 Å in respect to the corresponding residue in the PGP−bound state to avoid steric hindrance with the cytosine group of CDP-DAG. Meanwhile, the side chain of Arg65 moves toward the α-phosphoryl moiety of CDP-DAG due to electrostatic attraction (Figure 5B). Likewise, the residues involved in binding the G3P head group of PGP adopt different rotameric conformations compared to those in the CDP-DAG−bound state (Figure 5C). Specifically, Lys83, Arg110, Arg118, Lys137 and Tyr181 from one protomer move toward the 3’-phosphoryl moiety of PGP due to electrostatic attraction, while Arg110’ and Tyr181’ in the adjacent protomer move slightly away from it to avoid steric hindrance. The TM4-5 loop shifts toward TM6 by 1-3 Å so that the substrate-binding cavity opens wider in comparison with the CDP-DAG−bounds state (Figure 5A). On the intracellular surface of *Sa*PgsA−CDP-DAG complex structure, two channels (Channel 1 and Channel 2) connect the active site with cytoplasm and are separated by Lys75 from TM3 and Ala130 from TM4-5 loop (Figure 3B). While Channel 1 may serve as the portal for CMP release, Channel 2 likely guides the entry of G3P to the active site. In the PGP−bound state, the TM4-5 loop moves toward TM6, expanding Channel 1 at the cost of diminishing Channel 2. On the other hand, binding of CDP-DAG to the active site may induce conformational changes leading to the formation of Channel 2 to embrace G3P. Such reversible seesaw-like movement may facilitate the substrate entry and product release cycle during catalysis.

**Figure 5.**
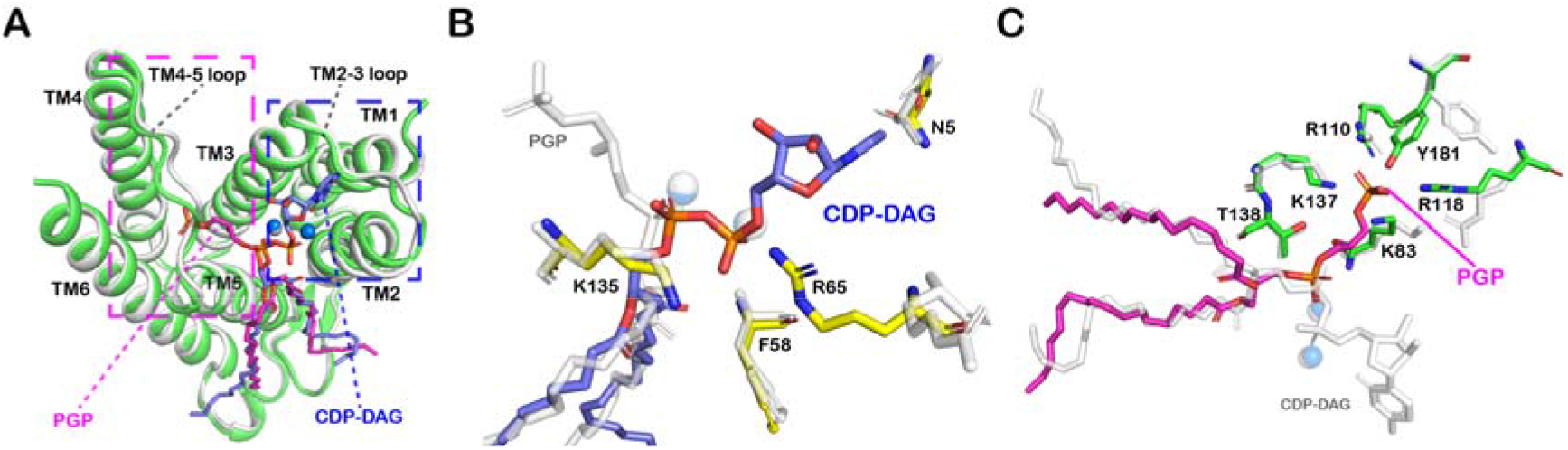
Structural dynamics of *Sa*PgsA in response to CDP-DAG or PGP binding. (**A**) Local conformational differences between the structures of *Sa*PgsA−CDP-DAG complex (white) and the *Sa*PgsA−PGP complex (green). The blue dashed box indicates conformational changes induced by CDP-DAG binding. The magenta dashed box indicates the conformational changes induced by the binding of PGP head group. PGP and CDP-DAG are presented as stick models in magenta and blue, respectively. (**B**) A zoom-in view of the conformation changes of amino acid residue side chains induced by CDP-DAG binding. The amino acid residues involved in binding the head group of CDP-DAG are presented as stick models in yellow (carbon), blue (nitrogen) and red (oxygen). For comparison, the reference structure (*Sa*PgsA−PGP complex) superposed on the one in complex with CDP-DAG is shown in silver. (**C**) An expanded view of the side chain conformation changes induced by PGP binding. The amino acid residues involved in binding the head group of PGP are presented as stick models in green (carbon), blue (nitrogen) and red (oxygen). As a reference, the structure of *Sa*PgsA−CDP-DAG superposed on the one in complex with PGP is shown in silver.

### A possible trade-off mechanism for PgsA-related daptomycin resistance

Due to misuse of antibiotics, development of antibiotic-resistance in *S. aureus* poses a global threat to human health [46]. Unfortunately, daptomycin treatment of MRSA infections can fail in more than 20% of total clinical cases due to the occurrence of daptomycin resistance [47]. Previous studies reported frequent occurrence of *pgsA* mutations in clinical and laboratory derived daptomycin-resistant *S. aureus* [18, 19, 25], *Streptococcus oralis* [37] and *Staphylococcus capitis* [24] strains. To further explore the mechanism of PgsA’s involvement in daptomycin resistance of Gram-positive pathogen, we constructed eight *Sa*PgsA single mutants which are directly from the daptomycin-resistant *S. aureus* or from *Streptococcus oralis* and *Staphylococcus capitis* strains [20, 24].

Among the eight *Sa*PgsA mutants (V59D, V59N, G61S, A64V, K75N, K135E, S177F and D187E) tested, all except A64V and D187E exhibited reduced activity when compared to the wild type (Figure 6A). As Asp187 is distant from the active site or the putative substrate entrance channels, the D187E mutation has little effect on the activity. In contrast, Val59, Gly61 and Ala64 are all located on TM2 nearby the characteristic and conserved helix kink essential for catalysis. In addition, Val59, Gly61 and Ala64 in the conserved “D_57_xxD_60_G_61_xxA_64_R_65_” motif are proximal to the bimetal catalytic center (Figure 6B). Therefore, the V59D and G61S mutations result in dramatic or complete loss of activity. As for V59N and A64V, they exhibit activity close to the wild type in detergent micelle, but show much lower activity in LCP. The different amphipathic environments may have dramatic influence on the binding site of CDP-DAG. Lys75 is located at TM3 near the strictly conserved residues Gly74 and Asp78 as well as the bimetal catalytic center. K75N mutation resulted in ∼50% loss of activity (Figure 6A). Presumably, the Asn75 residue introduced by mutation might form hydrogen bond with the hydroxyl groups of the ribose and distort/dislocate the substrate. Such mutation may also affect the release of PGP as it is located nearby the 3-phosphoryl moiety of PGP. Besides, Ser177 on TM6 is near the 3’-phosphoryl moiety of PGP. The S177F mutant also displayed dramatic reduction of activity. The mutation introduced a bulky side chain potentially causing severe clashes with PGP or G3P.

**Figure 6.**
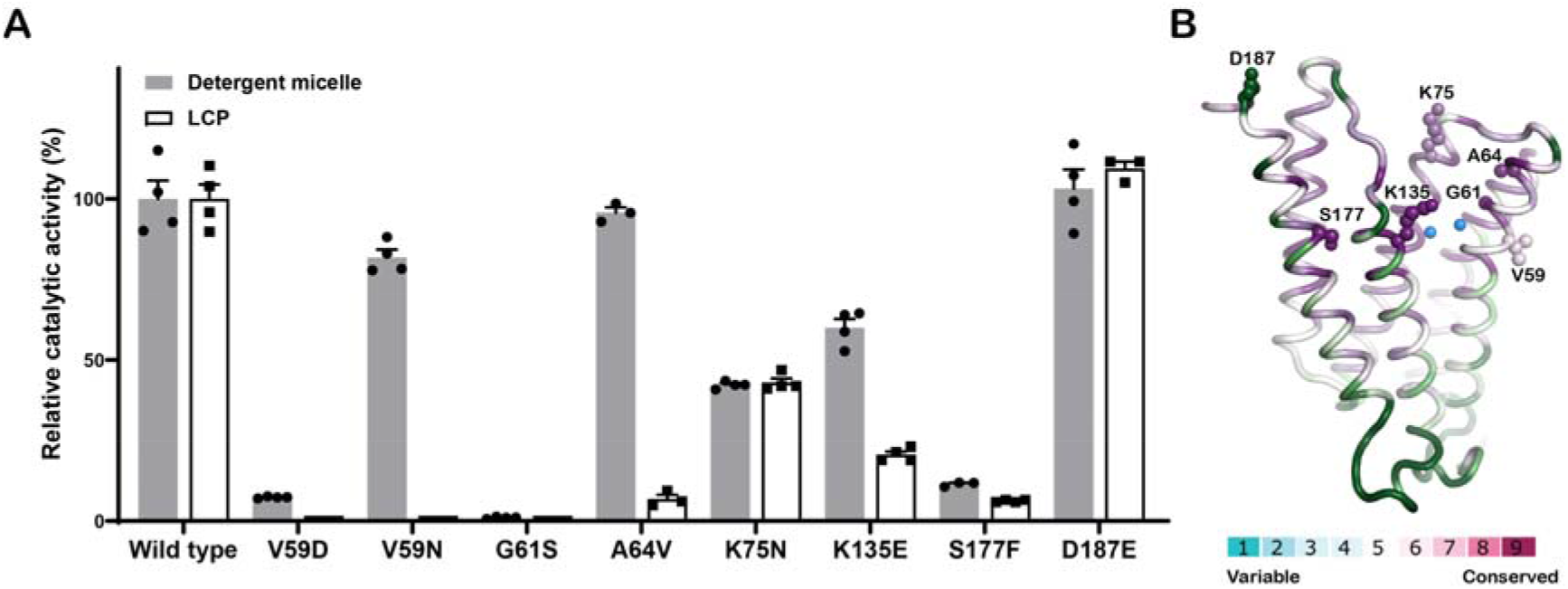
The daptomycin-resistant point mutants of *Sa*PgsA. (**A**) The relative enzymatic activity of the daptomycin-resistant point mutants. The error bars indicate ± SEM with n = 3 or 4. The solid grey bars represent the activity data measured with WT and mutant enzymes solubilized in detergent, whereas the white bars represent the data measured with enzymes embedded in LCP. (**B**) Structural mapping of the daptomycin-resistant mutations in *Sa*PgsA. The residues are colored on the basis of the conservation score of each residue. The amino acid residues subject to mutation for activity assay are shown as sphere models. For clarity, only one protomer of the *Sa*PgsA dimer is shown.

Collectively, the dysfunctional mutations of *Sa*PgsA might result in a significant decrease of PG content in *S. aureus* membrane. A recent study indicates that PG is involved in forming a tripartite complex with Ca^2+^-daptomycin and undecaprenyl-coupled cell envelope precursors, so as to support the action of daptomycin on bacterial cells by targeting and interrupting cell wall biosynthesis [48]. In case that PG content is lowered in the mutant strains, daptomycin might not be able to target cell wall biosynthesis, as it could not recruit enough PG to form the tripartite complex. As a result, the *S. aureus* strains carrying the *pgsA* mutations with null or low activity develop increased resistance (or reduced susceptibility) to daptomycin. Reduction of PgsA activity might effectively attenuate the bactericidal activity of daptomycin even though it would cause slower cell growth. Thus, *S. aureus* may trade-off PG biosynthesis with drug resistance to survive the antibiotic treatment while keeping anionic phospholipid content at minimal level.

## Discussion

PG, CL and their derivatives have pivotal functions in various biological processes within prokaryotic and eukaryotic cells [5, 6, 49], in addition to their fundamental roles as the building blocks of biological membranes. Back in 1960s, the activity of PgsA was discovered and the enzyme was partially purified from *E. coli* [42], and the bacterial *pgsA* gene was cloned in 1980s [50]. Further enzymatic kinetic study on *E. coli* PgsA suggested the enzyme adopts an order sequential Bi-Bi mechanism and CDP-DAG at high concentration may form a dead-end complex with the enzyme so as to inhibit the catalytic process [41]. Notably, *Sa*PgsA has recently been identified as a promising antibacterial target because of its vital role in the formation of bacterial cell membrane and cell wall [26]. Basing on the preliminary molecular docking result, we propose that the cajaninstilbene-acid analogue may specifically occupy the CDP-DAG binding site in *Sa*PgsA and lead to inhibition of its activity (Supplementary Fig. 8) [26]. Since CDP-DAG was previously found to inhibit PgsA activity by forming the so-called dead-end complex [41], the structures of *Sa*PgsA with CDP-DAG bound may provide useful insights to inspire the design and development of PgsA inhibitors as antibiotic drugs. Moreover, as the products of enzymes often serve to inhibit the activity [51], the structure of *Sa*PgsA−PGP complex may offer a different angle for the design of inhibitors.

Basing on the structural observation and functional analysis results, we propose a five-state working model and a putative nucleophilic substitution mechanism to account for the dynamic catalytic process mediated by PgsA (Figure 7 and Supplementary Fig. 9). Firstly, CDP-DAG from the membrane enters the active site through a lateral portal. Upon CDP-DAG binding, the protein may go through a conformational change around the membrane-perpendicular portal near the intracellular surface. The portal rearranges into two channels (Channel 1 and Channel 2) and the entry of G3P is guided by Channel 2, while Channel 1 is occupied by the head group of CDP-DAG. Secondly, the β-phosphorus of CDP-DAG (as an electrophile) and the 1-OH of G3P (as a nucleophile) are activated by the bimetal center and the overall acidic microenvironment of the active-site center. Zn1 serves to pull electrons from the β-phosphorus group towards the coordinating oxygen, making the β-phosphorus more electrophilic. When G3P is steered into the active site with its 1-OH in line with the di-phosphoryl of CDP-DAG, the nucleophilic attack launched by the 1-OH of G3P initiates a series of electron rearrangements and leads to the formation of new P-O bond between G3P and the β-phosphoryl of CDP-DAG (Supplementary Fig. 9). Thirdly, as the reaction proceeds, the products (PGP and CMP) form and CMP is released to cytosol through the expanded Channel 1. To further free up the active site, PGP diffuses outward to the bulk membrane through the same lateral portal used for CDP-DAG entry (Figure 7).

**Figure 7.**
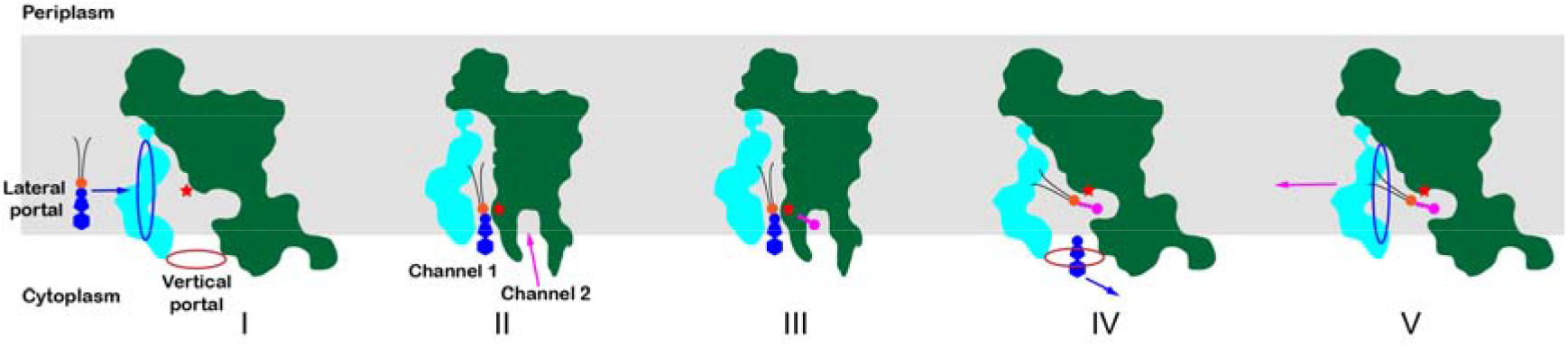
A multi-state cartoon model accounting for the catalytic process mediated by PgsA. The grey shading indicates the estimated membrane region. The red and blue elliptical rings indicate the vertical and lateral portals for the entry of G3P and CDP-DAG (or for the release of CMP and PGP) respectively. Among the five putative states (I-V), State II represents the one observed in the *Sa*PgsA−CDP-DAG complex structure, whereas State V corresponds to the state of the *Sa*PgsA−PGP complex structure.

Eukaryotic cardiolipin synthase catalyzes the conversion of CDP-DAG and PG to cardiolipin and CMP in mitochondria (Supplementary Fig. 10A) [52]. It is closely related to PgsA and they belong to the same family in the CDP-OH_P_transf superfamily (Supplementary Fig. 3A). Abnormalities in mitochondrial CL content are associated with several human diseases, such as Barth syndrome, myocardial ischemia/reperfusion injury, diabetes and Parkinson’s disease [53]. Besides, knockdown of cardiolipin synthase induces mitochondrial elongation in human cells [54]. While human cardiolipin synthase shares 29% sequence identity with *Sa*PgsA, the amino acid residues around the CDP-DAG and PGP binding sites in *Sa*PgsA are mostly conserved in human cardiolipin synthase (Supplementary Fig. 10B and C). As the structures of eukaryotic cardiolipin synthases are still unknown, the *Sa*PgsA structure can potentially serve as a homologous model for future researches on mammalian cardiolipin synthases.

## Materials and methods

### Cloning, expression and purification

The gene encoding *Sa*PgsA was synthesized (GenScript, China) with optimized codon usage for protein expression in *E. coli*. The target sequence was ligated into pET-15b vector between the NdeI and BamHI restriction enzyme sites. All point mutants of *Sa*PgsA were introduced through QuickChange site-directed mutagenesis and verified by sequencing. Sequences of all the primers used in this study are listed in Supplementary Table 2. Plasmids were transformed into *E. coli* C41(DE3) or C43(DE3) competent cells, then overexpressed in Terrific Broth medium at 37^℃^C. Once the cell density reached to OD_600_ of 1.0, the inducer isopropyl-β-D-1-thiogalactopyranoside (IPTG) was added with a final concentration of 0.5 mM. The induction of wild type and point mutant was continued for 2-3 h at 37 ℃ and 12-15 h at 16 ℃, respectively. Cells were harvested by centrifugation at 9,000×g for 15 min at 4 ℃ and stored at -80 ℃ refrigerator.

Protein purification procedures were performed at 4 °C. 50 g frozen cell paste was resuspended in 400 ml lysis buffer (50 mM Tris-HCl, 200 mM NaCl, 20 mM imidazole, pH 8.0) and then homogenized by using the T10 basic homogenizer (IKA, Germany). Cells were broken by passed through a French press (ATS Engineering, Canada) at pressure of 900-1000 bar for 4-5 times. The cell lysate was centrifuged at 12,000×g for 20 min to remove cell debris. The supernatant was centrifuged further at 160,000×g for 1 h to yield membrane. The membrane fraction was resuspended and homogenized in 100 ml lysis buffer, then solubilized with 1.5% (w/v) of n-dodecyl-β-D-maltopyranoside (β-DDM, Anatrace, USA). After solubilization for 2-3 h, insoluble materials were removed by centrifugation at 41,000×g for 30 min. The supernatant was then incubated with 3 ml pre-equilibrated Ni-NTA resin (GE Healthcare, USA).

After 2-3 h gentle agitation, the resin was packed to a gravity column and washed sequentially with 15 ml buffer A (25 mM Tris-HCl, 300 mM NaCl, 10% glycerin, 20 mM imidazole, 0.1% β-DDM, pH 7.5) and 15 ml buffer B (25 mM Tris-HCl, 300 mM NaCl, 10% glycerin, 70 mM imidazole, 0.05% β-DDM, pH 7.5). The protein was eluted with 12 ml buffer C (25 mM Tris-HCl, 300 mM NaCl, 10% glycerin, 300 mM imidazole, 0.05% β-DDM, pH 7.5), and immediately diluted by 2-folds volumes of buffer D (25 mM Tris-HCl, 300 mM NaCl, 10% glycerin, 0.05% β-DDM, pH 7.5). The protein sample was concentrated to 10-15 mg ml^-1^ (approximated by A_280_ absorbance) using a 50 kDa cut-off concentrator (Millipore, USA) and further purified by size exclusion chromatography (SEC) on a Superdex 200 increase 10/300 GL column (GE Healthcare, USA) in buffer E (10 mM Tris-HCl, 300 mM NaCl, 0.05% β-DDM, pH 7.5). Sharp monodisperse peak protein fractions of SEC were pooled and concentrated to 40-45 mg ml^-1^. And then, the protein samples were flash frozen in liquid nitrogen and stored at -80°C refrigerator.

### Crystallization

Crystals of *Sa*PgsA were grown in lipidic cubic phase (LCP) at 4°C or 20°C [31]. Initially, the protein solution was homogenized with 1.5-folds volumes molten 9.9 MAG (Molecular Dimensions, USA) using a gas-tight coupled syringe device (Hampton Research, USA) at room temperature (20-22°C). Crystallization trials were set up by covering 40 nl of LCP bolus with 800 nl of precipitant solution onto a 96-well glass sandwich plate (FAstal BioTech, China) using a Gryphon robot (Art Robbins Instruments, USA). The glass sandwich plate was immediately sealed with a glass coverslip, then stored and imaged in Rock Imager 1000 (Formulatrix, USA) at 20°C. Extensive crystallization trials yielded needle-like crystals, which were grown in 10-40 % (v/v) 1,4-butanediol, 0.2 M or 0.3 M zinc acetate, and 0.1 M imidazole/HCl (pH 5.0-8.5). Plate-like crystals mostly appeared in 20-25 days and matured to 100×100×20 μm about 40-60 days at 4°C. Meanwhile, we also obtained substrate-bound crystals in the crystallization condition containing CDP-DAG (Avanti, USA) and G3P (Sigma, USA). As for CDP-DAG, the amount of CDP-DAG was calculated on the basis of 10-folds protein mole number. The 9.9 MAG doped with CDP-DAG was prepared before it was used, and involved dissolving CDP-DAG in chloroform, adding it to molten 9.9 MAG in the appropriate amount, vortexing, and evaporating the chloroform with nitrogen gas for 3-4 min, then dried by a vacuum desiccator for 6-8 h. On the other hand, the protein solution was incubated with about 40-folds (molar ratio) G3P at 4°C for 1-2 h. The substrate-bound crystals appeared after 10 days and reached full size in 30 to 60 days at 20°C. Desired wells were opened with a glass cutter (Hampton Research, USA) and the crystals were harvested using 50-100 μm Dual Thickness Microloops LD (MiTeGen, USA) at 4°C or 20°C, respectively. The crystals were immediately flash frozen in liquid nitrogen without additional cryoprotection.

### Data collection and processing

X-ray diffraction data was collected at beamline BL18U1 of Shanghai Synchrotron Radiation Facility (SSRF, China) in the National Center for Protein Sciences Shanghai (NCPSS, China), using a PILATUS3 S 6M Detector (X-ray wavelength 0.9785 Å or 0.9793 Å). The near-Zn absorption K-edge anomalous data was collected at SPring-8 (Hyogo, Japan) beamline BL41XU, using an EIGER X 16M detector (X-ray wavelength 1.2820 Å). Diffraction images were indexed and integrated using XDS[55], scaled and merged with AIMLESS in the CCP4 software[56, 57]. The crystals appeared in the *C*2 space group, and diffracted X-ray to 2.5 Å (PGP−bound) and 3.0 Å (CDP-DAG−bound) resolution at synchrotron radiation source, respectively.

### Structure determination and refinement

Initial phases were obtained through *ab initio* phasing and chain tracing method using ARCIMBOLDO-LITE program with 10 copies of a helix containing 18 residues as an initial estimated models [32, 58]. Structure refinement was carried out through alternating between cycles of refinement in PHENIX (Refine) and manual building in Coot [59, 60]. Lipid, solvent and water molecules were added manually in the model for refinement at later stages when their electron densities were well identified. The *Sa*PgsA−PGP complex structure was refined to 2.5 Å resolution with *R*_work_/*R*_free_ values of 0.213/0.250, respectively. The *Sa*PgsA−CDP-DAG complex structure was solved by molecular replacement using the *Sa*PgsA−PGP complex structure as search model, and refined to 3.0 Å with *R*_work_/*R*_free_ values of 0.251/0.299. Data collection and refinement statistics are listed in Table S1. All structure figures were presentation with PyMOL [61]. The orientation and position of *S*aPgsA protein in the membrane was calculated by using the PPM web server [39]. The conservation score of each residue in *Sa*PgsA presented in Figure 6B was calculated by the ConSurf web server [62]. The omit maps for PGP (Supplementary Fig. 4A) and CDP-DAG (Supplementary Fig. 4C) were calculated using PHENIX (Polder) [63].

### Oligomeric state analysis through SEC-MALS

To analyze the oligomeric state of *Sa*PgsA in detergent micelle, the size-exclusion chromatography coupled with multi-angle light scattering (SEC-MALS) method was applied. 50 μl *Sa*PgsA sample was injected into the WTC-030S5 SEC column (Wyatt, USA) and eluted at 0.5 ml min^-1^ in buffer E. Data collection and analysis were performed with Astra 5 software (Wyatt, USA). The specific refractive index (dn dc^-1^) value of protein at 0.185 ml g^-1^ and that of β-DDM at 0.144 ml g^-1^ were used for data processing. The total molecular masses and individual masses of the proteins and the detergent were determined with Astra 5 software using protein conjugate analysis. The peak overlap and peak broadening corrections were carried out with Astra 5 software. Experiments were performed use protein sample with concentration at 1.13 mg ml^-1^, 4.54 mg ml^-1^ and 5.12 mg ml^-1^, respectively. All results showed that *Sa*PgsA was at dimeric state in detergent micelle. For clarity, only 4.54 mg ml^-1^ sample was presented in Supplementary Fig. 1D.

### Identification of PGP by mass spectrometry

To extract lipids, purified *Sa*PgsA (5.82 mg ml^-1^, 200 μl) was mixed with 40 mg SM2 Biobeads (Bio-rad, USA) and gently shaking overnight at 4 °C to remove detergent. The sample was centrifuged to remove detergent and Biobeads, then dried by a vacuum desiccator. A mixture of 60 μl chloroform, 120 μl methanol and 38 μl ultrapure water was added to the sample and then the tube was vortexed once every 2 min for 10 times. Then, a mixture of 60 μl chloroform and 60 μl of 2 M KCl was added and vortexed once every 2 min for 7 times. The mixture was centrifuged for 5 min at 18,800×g and the lower phase was separate out in a new tube. The selected lower phase sample was dried by a vacuum desiccator. Finally, add 25 μl chloroform and 25 μl methanol to the above tube, and the sample was ready for the following lipid mass spectrometry. The lipid mass spectrometry was performed according to the protocol described before [64]. Since there is no PGP mass spectrometry data in the existing database and the standard PGP sample is not commercially available, we manually searched for putative PGP molecule signal from total lipidomic data by identifying characteristic fragments peaks suggested from previous published data [15].

### Activity assay with radioisotope labeled substrate

*Sa*PgsA enzymatic activity was measured by the incorporation of [^14^C]-G3P into the chloroform-soluble PGP [42, 50]. A 50 μl reaction mixture contains 1.6 mM [^14^C]-G3P (specific activity at 129.3 mCi mmol^-1^, PerkinElmer, USA), 0.8 mM CDP-DAG (Avanti, USA), 100 mM Tris-HCl (pH 8.0), 100 mM MgCl2, 20 mM Triton X-100 (Sigma, USA) and 100 nM enzyme. The reaction system was incubated at 27 °C for 15 min. To terminate the reaction, 180 μl chloroform/methanol/concentrated HCl (1:2:0.02, v/v/v) was added, following by addition of 60 μl of chloroform and 60 μl of 2 M KCl per tube. The tube was vortexed for 1 min and centrifuged for 15 min at 2,400×g to separate the organic phase from the lower aqueous phase. Then pipette 100 μl lower organic phase to a new tube and centrifuged at 2,400×g for 15 min. For the measurement of chloroform-soluble PGP radioactivity, 40 μl lower organic phase was mixed with 1 ml OptiPhase Supermix Cocktail scintillation liquid (PerkinElmer, USA) and vortexed for 2 min. The sample was subjected to liquid scintillation counting in the Wallac 1450 MicroBeta trilux liquid scintillation and luminescence counter (PerkinElmer, USA). Three or four parallel repeat experiments were performed for the measurement of each data point.

In order to analyze the influence of divalent metal ions on the activity of *Sa*PgsA, 100 mM MgCl_2_ in the reaction buffer was replaced with 100 mM ZnCl_2_, BaCl_2_, CaCl_2_, CoCl_2_, MnCl_2_ or 10 mM EDTA, respectively. For the CDP-DAG- or G3P-dependent kinetic assays, 50 nM wild type *Sa*PgsA enzyme was used and the concentration of CDP-DAG/G3P was held constant at 0.8 mM/1.6 mM, whereas the concentration of the other substrate was varied from 0.025 to 1.6 or 0.025 to 0.8 mM as shown. For all the activity assays in detergent, G3P (≥95%, Cat. 94124, Sigma, USA) of high purity was used. The data were analyzed and plotted using GraphPad Prism 8.0.

### Activity assay in LCP

LCP was prepared using 9.9 MAG doped with CDP-DAG in different proportions as described above. Basing on the previous protocol [40], 2 μl sticky LCP was dispensed onto the sidewall of a UV-transparent 96-well plate (Cat. 8404, Thermo Scientific, USA). 200 μl pre-warmed buffer F (50 mM Tris-HCl, 0.1 mM EDTA, G3P and divalent metal ion at various concentration, pH 8.0) was loaded to each well. The reaction was immediately monitored in a plate reader with 271 nm absorbance at 30°C. Using CMP standards prepared as stock solutions in 0.1 N HCl [65], the molar extinction coefficient of CMP in the assay buffer was calibrated as 9,175 M^-1^ cm^-1^. For CDP-DAG *K*_m_ determination: 50 μg ml^-1^ of *Sa*PgsA, 0-3.5 mol% CDP-DAG, 1 mM G3P (Cat. 94124, Sigma, USA) and 0.1 M MgCl_2_ were used. For G3P *K*_m_ determination: 50 μg ml^-1^ of *Sa*PgsA, 2 mol% CDP-DAG, 0-10 mM G3P and 0.1 M MgCl_2_ were used. For the activity assays of single point mutants: 100 μg ml^-1^ of enzyme, 0.5 mol% CDP-DAG, 15 mM glycerol phosphate isomeric mixture (Cat. G6501, Sigma, USA) and 0.1 M MgCl_2_ were used. For effect of divalent metal ions on *Sa*PgsA activity: 50 μg ml^-1^ *Sa*PgsA, 0.5 mol% CDP-DAG, 15 mM glycerol phosphate isomeric mixture and various metals used as follows (MgCl_2_, 0.1 M; ZnCl_2_, 0.63 mM; MnCl_2_, 0.4 mM; CoCl_2_, 12.5 mM; CdCl_2_, 2.5 mM). The glycerol phosphate isomeric mixture contains the β-isomer (G2P) and the racemic α-isomers (G1P and G3P). While High purity G3P (≥95%) was used for the assays of [CDP-DAG] and [G3P] kinetics in LCP, the glycerol phosphate isomeric mixture at high concentrations (15 mM) was used for other activity assay in LCP to reduce cost, as we found that G1P and G2P did not inhibit *Sa*PgsA activity even at 10-fold molar excess of G3P. For measuring *K*m values, three independent experiments were carried out, each with three such technical replicates. Other assays were performed with three or four technical replicates where the same batch of LCP was dispensed into three or four wells. The data were analyzed and plotted using GraphPad Prism 8.0.

## Data availability

The atomic models and structure factors have been deposited with the Protein Data Bank under the accession number 7DRJ and 7DRK. Enzymatic activity assay data presented in Figures 2B, 3E, 3F, 4A, 4B, 4E, 6A, S5B and S5C are accessible through http://dx.doi.org/10.17632/3d6w948vf5.1 webpage. There are no restrictions on data availability.

## Acknowledgements

We thank Dr. Isabel Usón at the Institute of Molecular Biology of Barcelona for the guidance in using the ARCIMBOLDO-LITE program and helpful discussion; the staff members at BL18U1 of Shanghai Synchrotron Radiation Facility (SSRF) and BL41XU of SPring-8 for their assistance during X-ray data collection; Y. Wang, H. J. Zhang, X.X. Yu and Z. S. Xie at the Protein Science Research Facility of the Institute of Biophysics (IBP) for their help in LCP crystallization, activity assay, static light-scattering experiments and mass spectrometry analysis respectively; Y. Yin for the efforts in protein expression and purification; X. B. Liang for technical assistance on biochemistry, crystal handling and X-ray diffraction data collection. The project is financially supported by the National Natural Science Foundation of China (31925024 and 31670749, Z.L.; 31870726, D.L.), the Strategic Priority Research Program of CAS (XDB37020101, Z.L.; XDB37020204, D.L.), the Basic Frontier Science Research Program of CAS (ZDBS-LY-SM003) and CAS Facility-based Open Research Program.

## Author Contributions

B.Y. cloned, expressed and purified the wild type and mutants of *Sa*PgsA, crystallized *Sa*PgsA, collected and processed X-ray diffraction data, solved and refined the structures, and carried out activity assay with radioisotope-labeled substrate. H.Y. performed *Sa*PgsA activity assay in LCP under D.L.’s supervision. B.Y., D.L and Z.L. analyzed the structures and catalytic mechanism. B.Y. and Z.L. wrote the initial draft and all authors participated in revising the manuscript. Z.L conceived and supervised the project.

