## Supplementary Figures 1-10 and Tables 1-2 for "The phosphatidylglycerol phosphate synthase PgsA utilizes a trifurcated amphipathic cavity for catalysis at the membrane-cytosol interface"

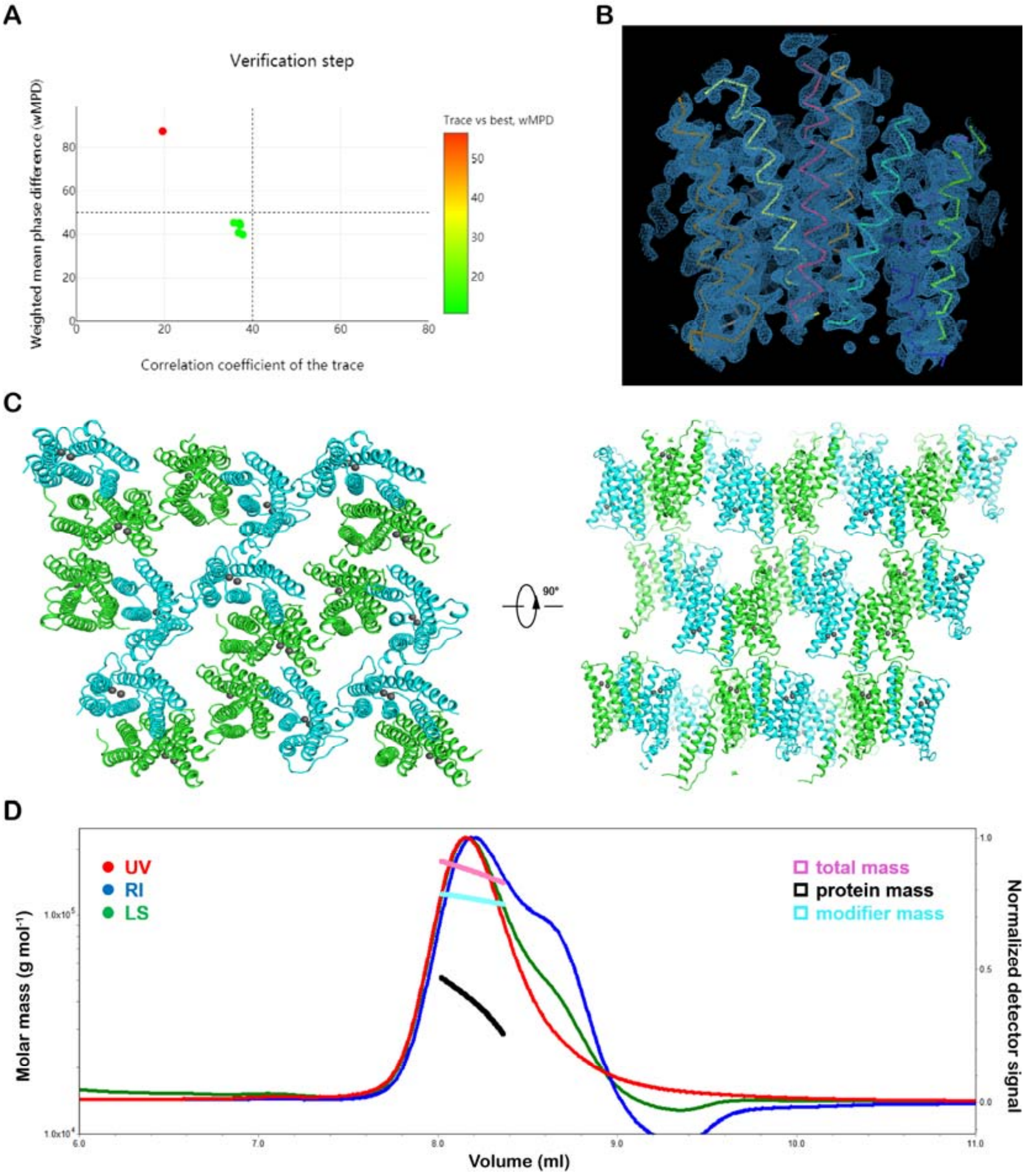

**Supplementary Figure 1. Structure determination, crystal packing and oligomeric state analysis of *Sa*PgsA.**

(**A**) Verification step of structure determination through the ARCIMBOLDO-LITE program**.** The verification result indicates that the structure is solved as the best solution (red dot) is clearly distinguished from a random one (green dots). The "coil coiled mode" in ARCIMBOLDO-LITE incorporates an additional verification step that generates perturbations of the substructure leading to the best solution and compares their scores before and after extension [1].

(**B**) The $2F_o$–$F_c$ electron density map calculated with the output model of ARCIMBOLDO-LITE program. The contour level is at $1.5 \times \sigma$ level.

(**C**) Packing of the *Sa*PgsA dimers in the crystal lattice. Two-dimensional lamina are formed by packing of *Sa*PgsA dimers and inverted ones along the planes parallel to the *b* and *c* axes, and the 2D lamina further stack along the direction of *a* axis in three-dimensional space to form the type I membrane protein crystal [2]. Two orthogonal views of the crystal lattice are shown**.**

(**D**) The oligomeric state of *Sa*PgsA in detergent solution. The UV absorbance, refractive index (RI) and laser light-scattering (LS) profiles of *Sa*PgsA eluted in β-DDM solution were measured through the size-exclusion chromatography with multi-angle light scattering (SEC-MALS) method. The total mass of *Sa*PgsA protein-detergent micelle complex is estimated at $1.580\times10^5$ (±0.5%) Da, while the measured protein mass is at $4.005\times10^4$ (±0.5%) Da, close to the theoretical mass of the *Sa*PgsA dimer ($4.635\times10^4$ Da).

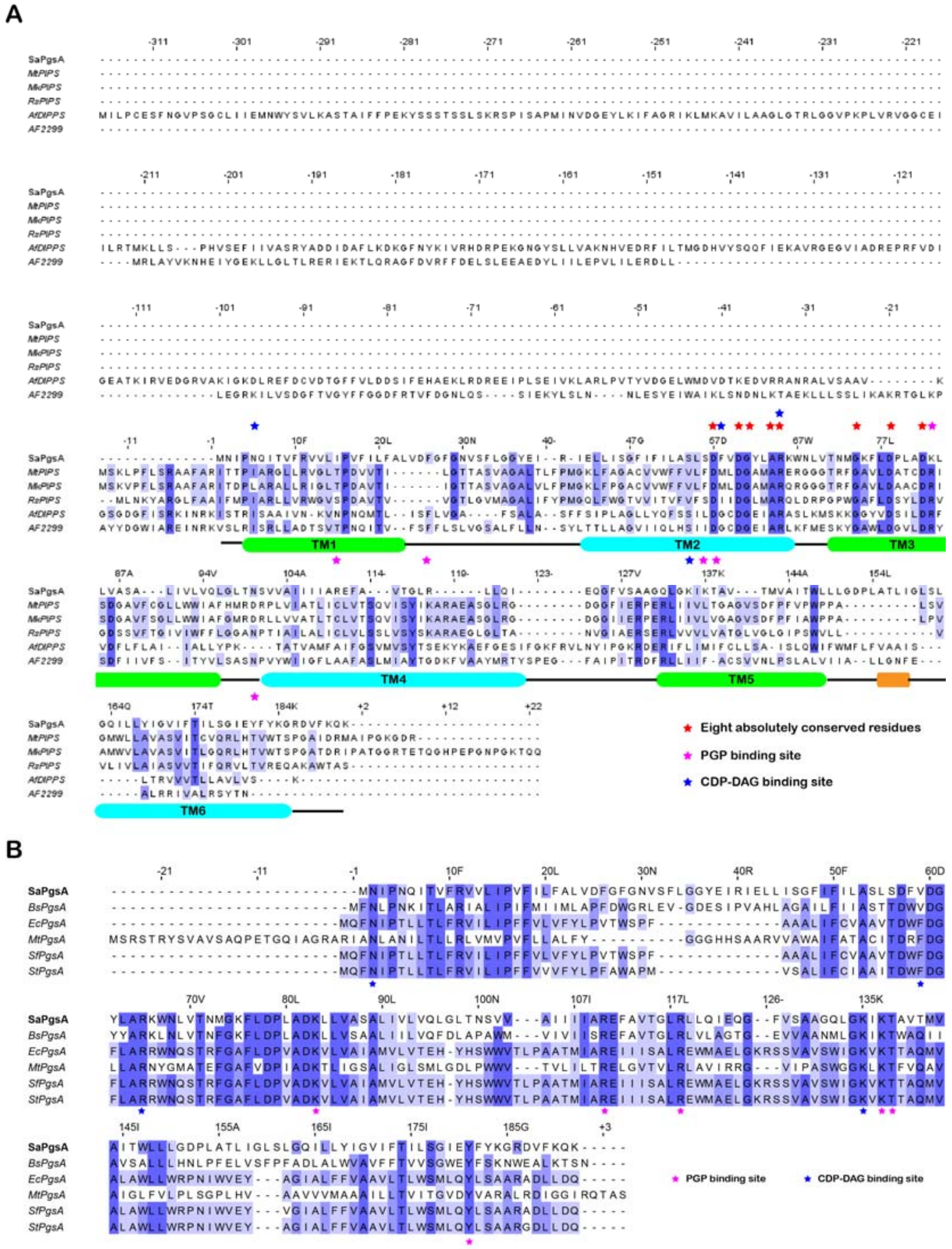
A
SaPgsA
MtPIPS
MkPIPS
RsPIPS
AfDIPPS
AF2299
TM1
TM2
TM3
TM4
TM5
TM6
Eight absolutely conserved residues
PGP binding site
CDP-DAG binding site
B
SaPgsA
BsPgsA
EcPgsA
MtPgsA
SfPgsA
StPgsA
PGP binding site
CDP-DAG binding site

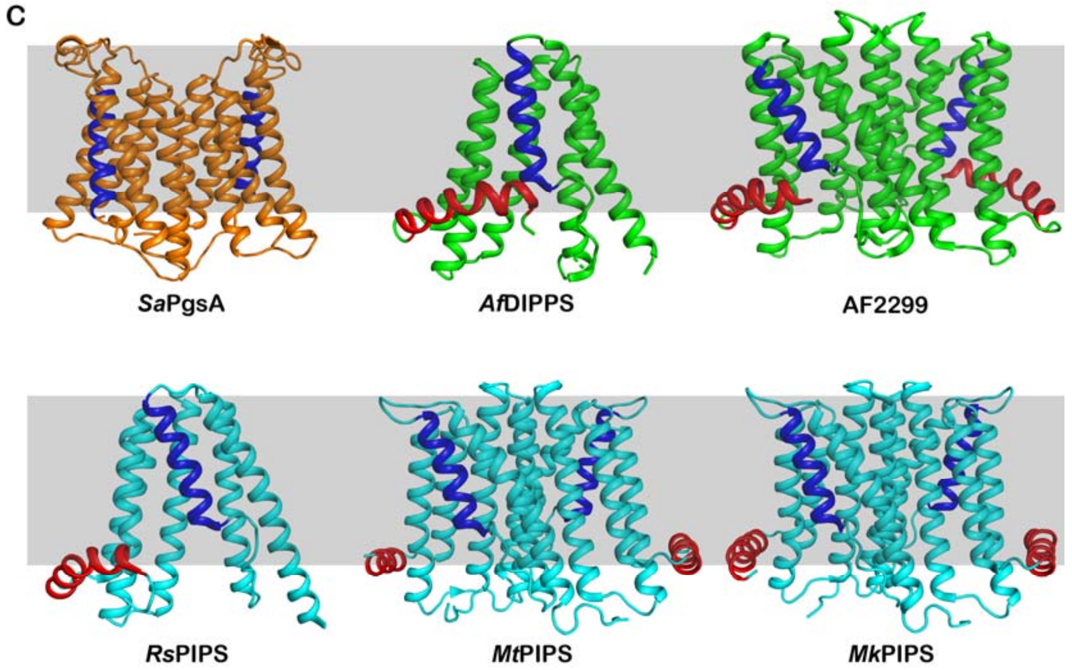

**Supplementary Figure 2. Comparison of *Sa*PgsA with other members of CDP-alcohol phosphotransferase.**

**(A** and **B)** Sequence alignment of *Sa*PgsA with the sequences of other known CDP-OH_P_transf structures (**A**) or prokaryotic PgsA homologs (**B**). The alignment was performed by using the Clustal Omega web server and presented using Jalview [3, 4]. Each of residue is coloured on the basis of their sequence percentage identity from white to blue. Secondary structural elements of *Sa*PgsA are marked on the alignment result. Abbreviation of species names: *Af, Archaeoglobus fulgidus; Bs, Bacillus subtilis; Ec, Escherichia coli; Mk, Mycobacterium kansasii; Mt, Mycobacterium tuberculosis; Rs, Renibacterium salmoninarum; Sf, Shigella flexneri; St, Salmonella typhimurium.*

(**C**) Comparison of *Sa*PgsA with the other members of CDP-alcohol phosphotransferase superfamily. PDB codes of *Af*DIPPS, AF2299, *Rs*PIPS, *Mt*PIPS and *Mk*PIPS are 4MND, 4O6M, 5D91, 6H59 and 6WMV, respectively. The characteristic long N-terminal juxta-membrane helix of all other known CDP-OH_P_transf structures is colored in red. The shorter TM5 is colored in blue. The approximate position of the membrane hydrophobic region is marked with grey shading.

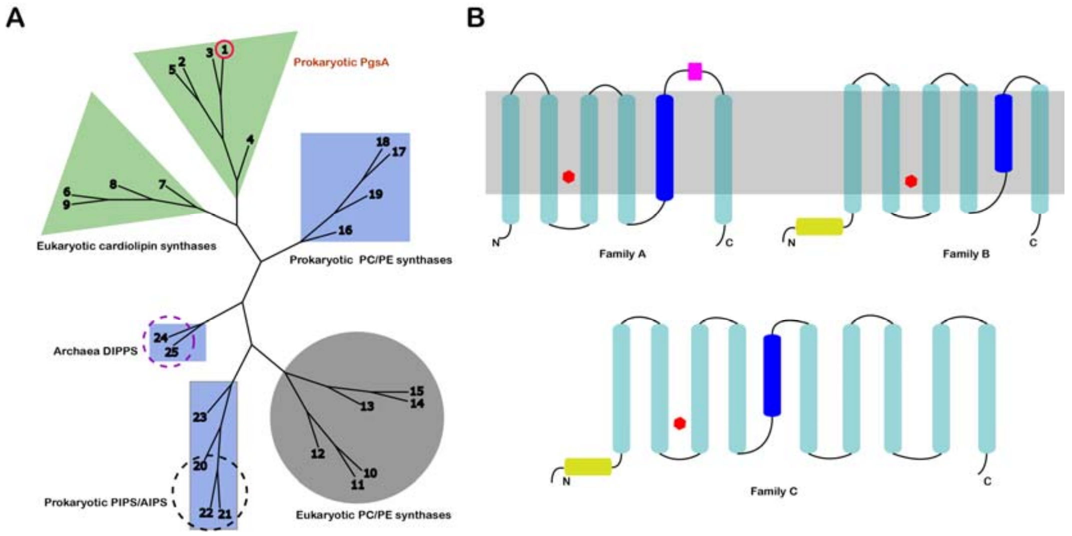

**Supplementary Figure 3. Phylogenetic tree and protein topology models for three families of CDP-OH_P_transf superfamily.**

(**A**) Phylogenetic tree of the three families of CDP-OH_P_transf superfamily. Multiple alignment of 25 sequences was performed using ClustalW method on MEGA X [5], then conducted unrooted phylogenetic analyses with neighbor joining (NJ) method. The visualization was achieved using T-rex [6] and manually edited for clarity. Triangle with green shading is indicating family A, rectangle with blue shading is indicating family B and circle with grey shading is indicating family C. The red circle highlights the target protein *Sa*PgsA, while the cyan star labels the cardiolipin synthase from human (*Hs*CLS); The purple dashed circle indicates two archaeal DIPPS with known structures (*Af*DIPPS and AF2299, PDB code: 4MND and 4O6M); the dark dashed circle labels three PIPS homologs with known structures, namely *Mt*PIPS, *Mk*PIPS and *Rs*PIPS (PDB code: 6H69, 6WMV and 5D91). The UniProt IDs of sequences used to generate the tree are as follows: (1) P63756; (2) P0ABF8; (3) P46322; (4) P9WPG3; (5) Q7CQB9; (6) Q9UJA2; (7) Q07560; (8) Q93YW7; (9) Q80ZM8; (10) Q8BGS7; (11) Q9Y6K0; (12) Q28H54; (13) Q8WUD6; (14) Q8C025;

(15) Q1LZE6; (16) D5KX81; (17) A9CIM3; (18) Q9KJY8; (19) Q98MN3; (20) A9WSF5; (21) P9WPG7; (22) U5WZP7; (23) O58215; (24) O29976; (25) O27985.

(**B**) Cartoon models showing the topologies of proteins belonging to three families of CDP-OH_P_transf superfamily. The α-helices are presented as cylinders. The active center formed by conserved sequence motif between TM2 and TM3 is represented by a red hexagon. The shortest helix TM5 is coloured in blue. The distinguishing additional helix between TM5 and TM6 of *Sa*PgsA is coloured in magenta. The characteristic long N-terminal juxta-membrane helix of family B and C is colored in yellow. The approximate position of the membrane-embedded hydrophobic region is marked with grey shading.

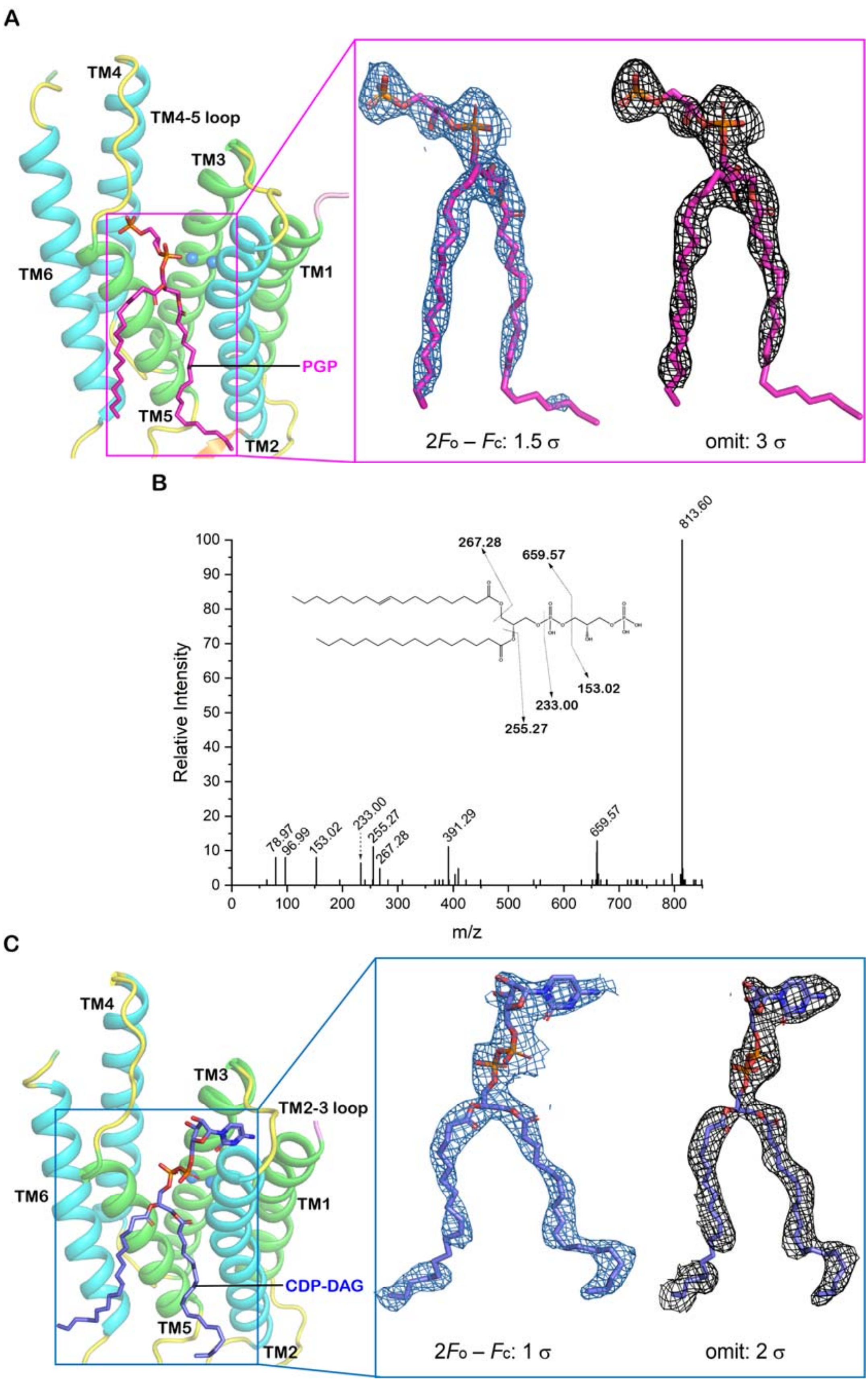

**Supplementary Figure 4. Validation of phospholipid molecules at active sites of *Sa*PgsA.**

(**A**) Electron density maps of PGP. The $2F_o$–$F_c$ (1.5 σ) and omit (3 σ) electron density maps for PGP are shown in inset. PGP is shown as magenta stick. Zinc ions are presented as marine blue spheres. The adjacent protomer is omitted for clarity.

(**B**) Mass spectrometry analysis of the lipid extracted from the *Sa*PgsA sample. The observed intact molecule and fragments peaks are consistent with the putative PGP and its fragment ions shown in the inset.

(**C**) Electron density maps of CDP-DAG. The $2F_o$–$F_c$ (1 σ) and omit (2 σ) electron density maps for CDP-DAG are shown in inset. CDP-DAG is shown as light blue stick. Zinc ions are presented as marine blue spheres. The adjacent protomer is omitted for clarity.

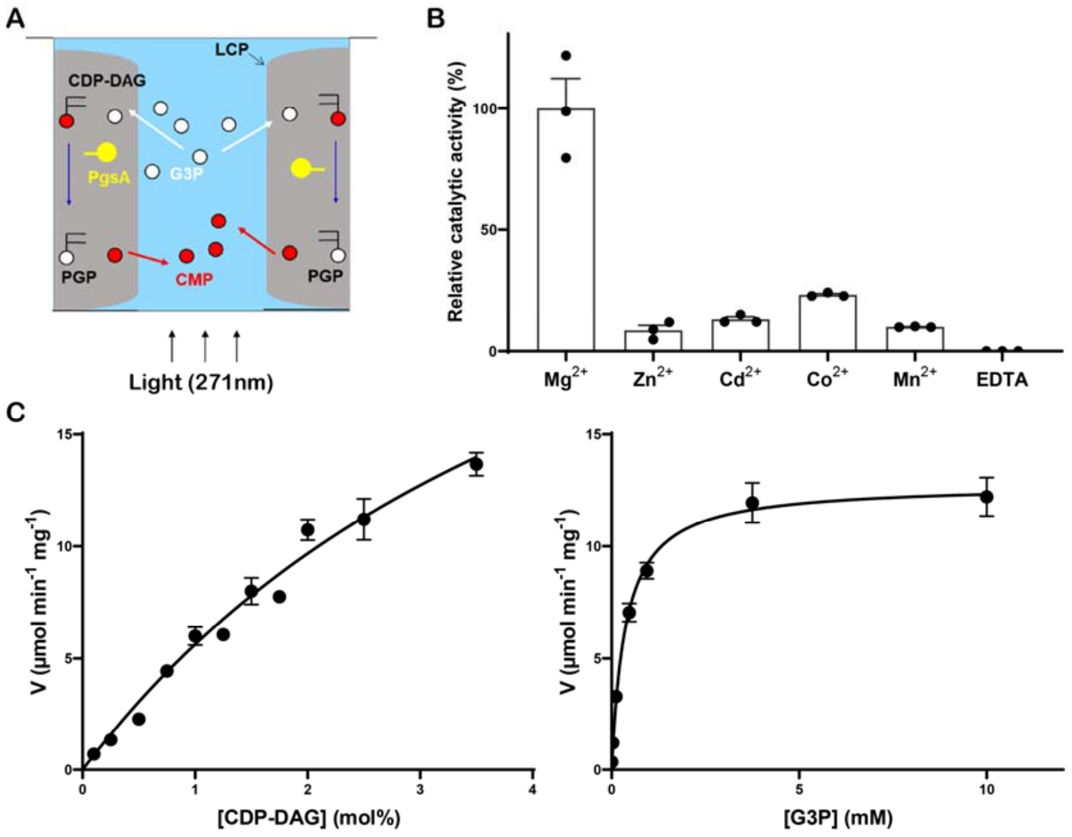

**Supplementary Figure 5. Activity assay of *Sa*PgsA in lipid cubic phase (LCP).**

(**A**) A cartoon diagram illustrating the principle of the assay of *Sa*PgsA activity in LCP. LCP (grey) reconstituted with the lipophilic CDP-DAG and *Sa*PgsA was deposited to the sidewall of 96-well plate. Because LCP is sticky, it stays where it is put and stays out of the light path (arrows at the bottom). Therefore, CDP-DAG and *Sa*PgsA, being confined in LCP, do not give UV absorbance signal. The assay is initiated by adding the buffer (blue) containing G3P and divalent metal ion (omitted in the figure for clarity). The water-soluble substrate G3P and divalent metal cofactor diffuse into the nanoporous LCP. The enzyme then catalyzes the reaction, generating the UV-absorbing CMP. Being water soluble, CMP is released to the soaking solution, into the light path where it is detected by a plate reader.

(**B**) Effect of divalent metal ions on *Sa*PgsA activity in LCP. The error bars indicate ± SEM with n = 3.

(**C**) Substrate-dependent kinetic measurement of *Sa*PgsA in LCP. The activity of *Sa*PgsA did not reach plateau with [CDP-DAG] at around 3.5 mol%. Because CDP-DAG at the concentration higher than 3.5 mol% begins to cause phase transition in LCP and thus makes the assay difficult, the activity of *Sa*PgsA with CDP-DAG concentrations higher than 3.5 mol% could not be measured accurately in LCP. With the current data measured in LCP, the $K_m$ value could be obtained for CDP-DAG. Average and standard error of the mean of three independent experiments (three parallel measurements per experiment) are reported.

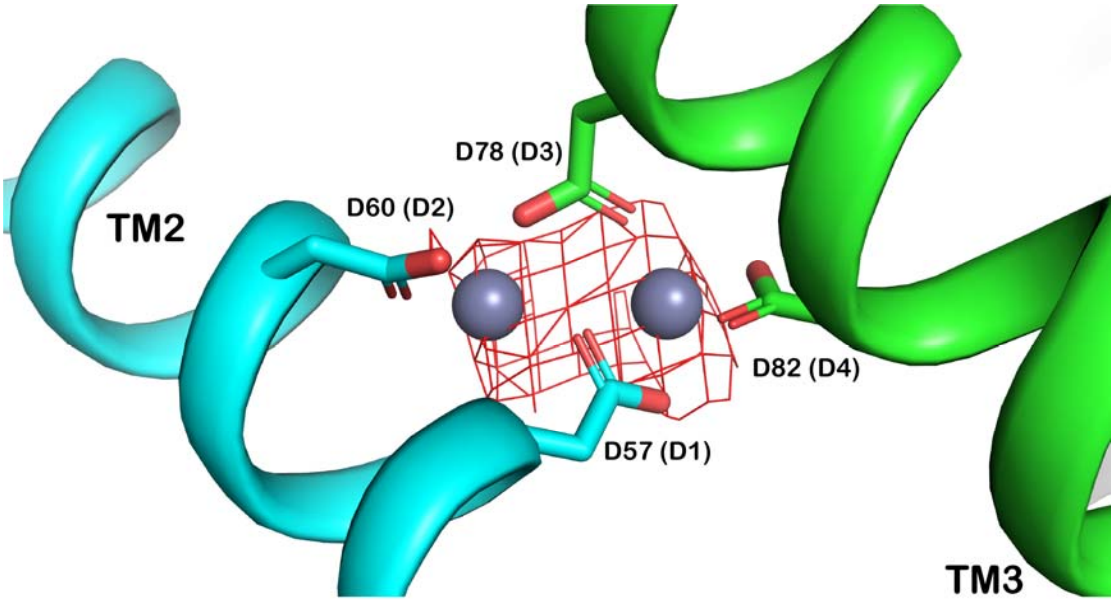

**Supplementary Figure 6. Validation of metal ions in the active site of *Sa*PgsA.**

The anomalous difference Fourier density map (3.0 σ) calculated from the dataset collected at a wavelength (1.282 Å) close to the K-edge of Zn. The zinc ions are shown as grey spheres. D1, D2, D3 and D4 within parentheses indicate the serial number corresponding to the conserved sequence motif (D1xxD2G1xxAR…G2xxxD3xxxD4).

**A**

| ID | Res. | Metal | Occupancy | B factor (env.)[1] | Ligands | Valence[2] | nVECSUM[3] | Geometry[1,4] | gRMSD(°)[1] | Vacancy[1] | Bidentate | Alt. metal |
|---|---|---|---|---|---|---|---|---|---|---|---|---|
| C:1 | ZN | Zn | 1 | 51.6 (51.9) | $O_6$ | 2.3 | 0.085 | Octahedral | 16.4° | 0 | 0 | Mg, Co, Ni, Mn, Zn, Cu |
| C:2 | ZN | Zn | 1 | 51.7 (53.4) | $O_5$ | 2.1 | 0.065 | Trigonal Bipyramidal | 9° | 0 | 0 | Co, Zn |
| C:3 | ZN | Zn | 1 | 48.2 (62) | $O_4$ | 1.5 | 0.22 | Tetrahedral | 16.2° | 0 | 0 | Cu |
| C:4 | ZN | Zn | 1 | 53.5 (50.3) | $O_6$ | 2.3 | 0.11 | Octahedral | 16.9° | 0 | 0 | Mg, Co, Ni, Mn, Zn, Cu |

Legend: Not applicable | Outlier | Borderline | Acceptable

**B**

| ID | Res. | Metal | Occupancy | B factor (env.)[1] | Ligands | Valence[2] | nVECSUM[3] | Geometry[1,4] | gRMSD(°)[1] | Vacancy[1] | Bidentate | Alt. metal |
|---|---|---|---|---|---|---|---|---|---|---|---|---|
| A:201 | ZN | Zn | 1 | 51.2 (54.7) | $O_6$ | 1.5 | 0.12 | Trigonal Bipyramidal | 2.3° | 0 | 1 | |
| A:301 | ZN | Zn | 1 | 59.7 (55.4) | $O_6$ | 1.5 | 0.11 | Tetrahedral | 16° | 0 | 2 | Cu |
| B:201 | ZN | Zn | 1 | 58.5 (55.4) | $O_6$ | 1.3 | 0.041 | Trigonal Bipyramidal | 18.1° | 0 | 1 | |
| B:301 | ZN | Zn | 1 | 43.7 (50.6) | $O_5$ | 1.4 | 0.15 | Trigonal Bipyramidal | 11.5° | 0 | 0 | |

Legend: Not applicable | Outlier | Borderline | Acceptable

**Supplementary Fig 7. Metal-ligand geometry and valence through the CheckMyMetal web server [7].**

(**A**) *Sa*PgsA–PGP complex structure.

(**B**) *Sa*PgsA–CDP-DAG complex structure.

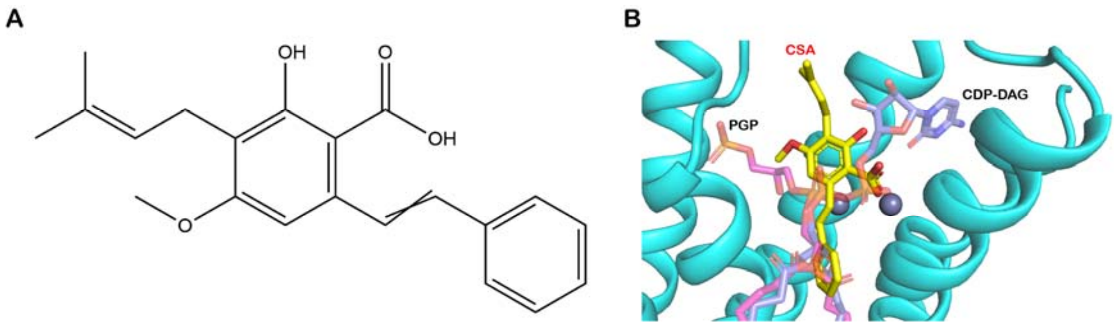

**Supplementary Fig 8. Molecular docking of cajaninstilbene-acid.**

(**A**) Structural formula of cajaninstilbene-acid (2-Hydroxy-4-Methoxy-3-Prenyl-6-Styrylbenzoic Acid).

(**B**) Binding site of cajaninstilbene-acid molecule in *Sa*PgsA suggested by molecular docking.

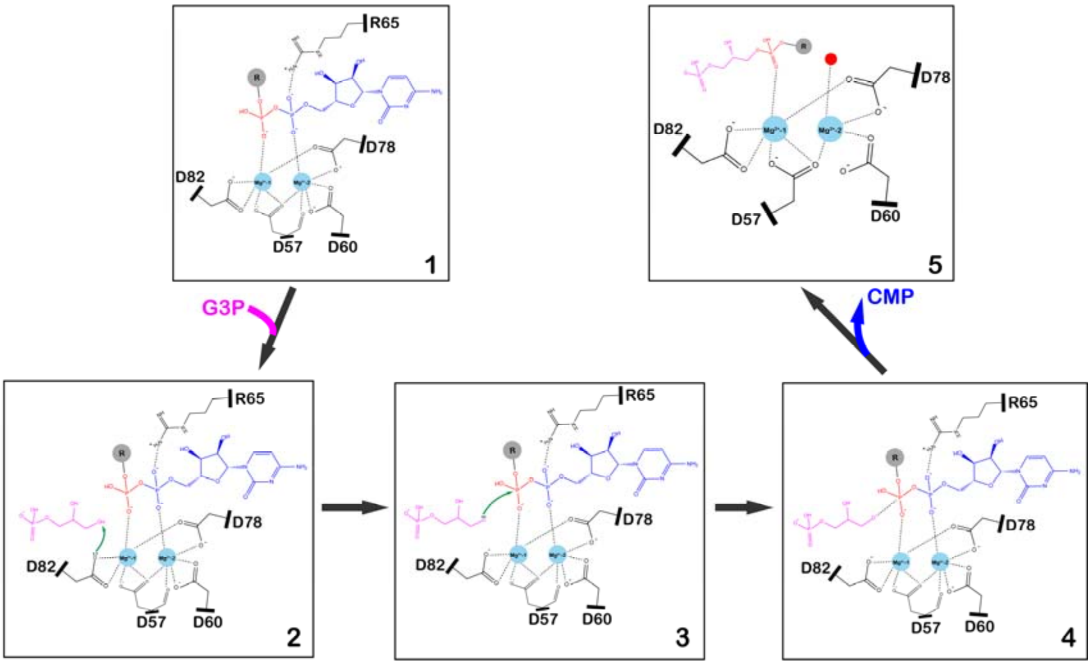

**Supplementary Fig 9. A nucleophilic substitution reaction model for PGP biosynthesis in the active site of *Sa*PgsA.**

State 1 is represented by the SaPgsA−CDP-DAG complex structure, States 2, 3 and 4 are defined as putative transition states, State 5 is indicated by the SaPgsA−PGP complex structure.

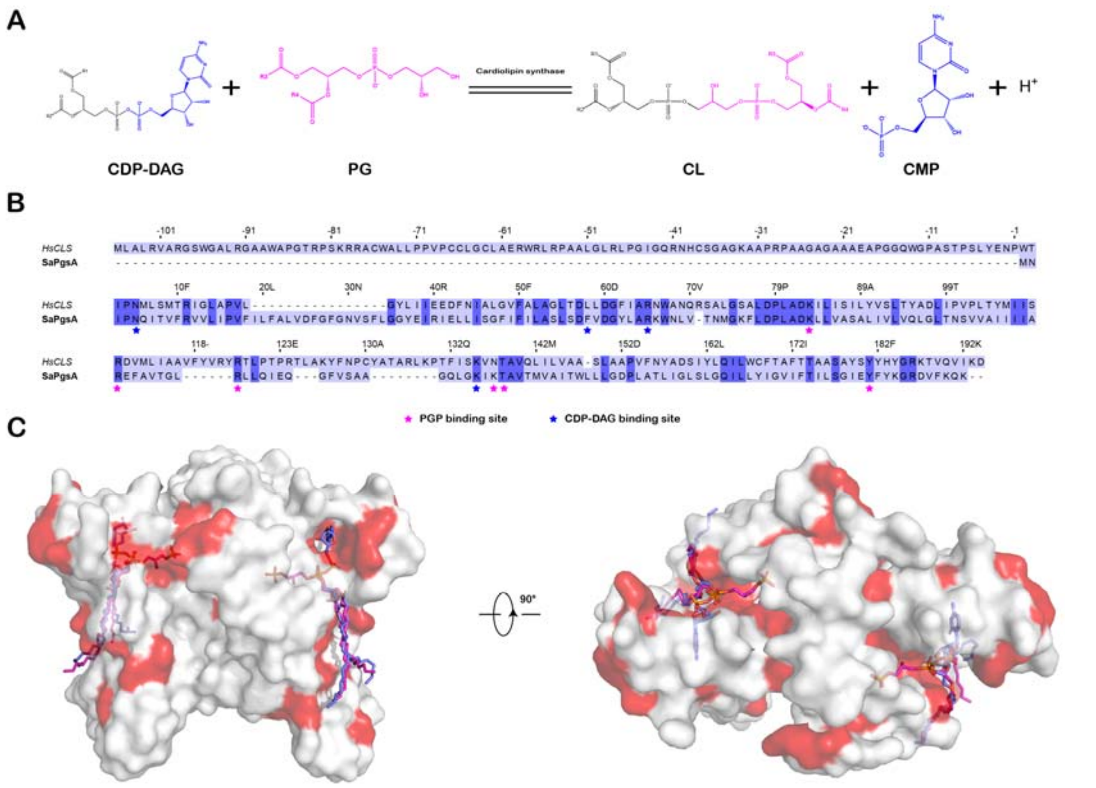

**Supplementary Fig 10. Close evolutionary relationship between PgsA and human cardiolipin synthase.**

(**A**) Cardiolipin synthase from the mitochondria of eukaryotic cells catalyzes the reaction of CDP-DAG with PG to generate CL and CMP.

(**B**) Alignment of the amino acid sequences of human cardiolipin synthase (*Hs*CLS) and *Sa*PgsA.

(**C**) The identical residues between *Hs*CLS and *Sa*PgsA is mapped on the *Sa*PgsA structure and highlighted in red.

**Supplementary Table 1. Data collection, processing and structure refinement statistics.**

| | ***Sa*PgsA–PGP (for phasing)** | ***Sa*PgsA–CDP-DAG** | ***Sa*PgsA /Zn-edge** |
|---|---|---|---|
| **Data collection** | | | |
| Space group | *C*2 | *C*2 | *C*2 |
| Cell dimensions | | | |
| *a*, *b*, *c* (Å) | 127.54, 59.38, 72.99 | 127.78, 59.43, 73.03 | 128.94, 59.28, 73.04 |
| *β* (°) | 106.18 | 106.66 | 106.15 |
| Wavelength (Å) | 0.9793 | 0.9785 | 1.2820 |
| Resolution (Å) | 29.69 - 2.50 (2.59 - 2.50) | 9.99 - 3.00 (3.10 - 3.00) | 45.23 – 3.99 (4.13 – 3.99) |
| $R_{merge}$ | 0.169 (1.510) | 0.171 (0.961) | 0.266 (0.687) |
| $R_{pim}$ | 0.074 (0.692) | 0.088 (0.487) | 0.136 (0.803) |
| $I/\sigma(I)$ | 8.44 (1.05) | 7.81 (1.61) | 7.02 (3.61) |
| Completeness (%) | 98.41 (96.67) | 96.35 (99.52) | 99.00 (96.70) |
| Redundancy | 5.9 (5.5) | 4.6 (4.7) | 13.5 (12.3) |
| $CC_{1/2}$ | 0.997 (0.458) | 0.994 (0.685) | 0.997 (0.951) |
| **Refinement** | | | |
| Resolution (Å) | 29.69 - 2.50 | 9.99 - 3.00 | |
| No. reflections | 18,027 | 10,317 | |
| $R_{work}$ | 0.213 (0.337) | 0.251 (0.284) | |
| $R_{free}$ | 0.250 (0.332) | 0.299 (0.389) | |
| No. atoms | | | |
| Protein | 2,899 | 2,835 | |
| Ligand/ion | 335 | 333 | |
| Water | 147 | 145 | |
| *B*-factors | | | |
| Protein | 54.70 | 54.38 | |
| Ligand/ion | 63.67 | 62.45 | |
| Water | 64.12 | 44.59 | |
| RMS deviations | | | |
| Bond lengths (Å) | 0.002 | 0.004 | |
| Bond angles (°) | 0.48 | 0.92 | |
| Ramachandran favored (%) | 99.19 | 98.08 | |
| Ramachandran allowed (%) | 0.81 | 1.92 | |
| Ramachandran outliers (%) | 0.00 | 0.00 | |
| PDB code | 7DRJ | 7DRK | |

Values in parentheses are for highest-resolution shell.

**Supplementary Table 2. Sequences of the primers used in the study of *Sa*PgsA.**

| Primer Name | Primer Sequence (5' to 3') |
|---|---|
| D57A_Forward | CTTCCCTTAGCGCGTTTGTTGATGG |
| D57A_Reverse | CCATCAACAAACGCGCTAAGGGAAG |
| D60A_Forward | CTTCCCTTAGCGATTTTGTTGCGGGTTATTTAGCTAGAAAATGG |
| D60A_Reverse | CCATTTTCTAGCTAAATAACCCGCAACAAAATCGCTAAGGGAAG |
| D78A_Forward | CAAATATGGGGAAATTTTTGGCGCCATTAGCGGATAA |
| D78A_Reverse | TTATCCGCTAATGGCGCCAAAAATTTCCCCATATTTG |
| D82A_Forward | GATCCATTAGCGGCGAAATTATTAGTTG |
| D82A_Reverse | CAACTAATAATTTCGCCGCTAATGGATC |
| K83A_Forward | GATCCATTAGCGGATGCGTTATTAGTTGCAAG |
| K83A_Reverse | CTTGCAACTAATAACGCATCCGCTAATGGATC |
| R110A_Forward | CATTATTATTGCCGCGGAATTTGCCGTAAC |
| R110A_Reverse | GTTACGGCAAATTCCGCGGCAATAATAATG |
| R118A_Forward | CCGTAACTGGTTTAGCGTTACTACAAATTG |
| R118A_Reverse | CAATTTGTAGTAACGCTAAACCAGTTACGG |
| K137A_Forward | CTGGTCAATTAGGTAAAATTGCGACAGCAGTTACTATGGTAGC |
| K137A_Reverse | GCTACCATAGTAACTGCTGTCGCAATTTTACCTAATTGACCAG |
| T138A_Forward | GTAAAATTAAAGCGGCAGTTACTATG |
| T138A_Reverse | CATAGTAACTGCCGCTTTAATTTTAC |
| Y181A_Forward | CTTATCTGGTATTGAAGCGTTTTATAAAGGTAGAG |
| Y181A_Reverse | CTCTACCTTTATAAAACGCTTCAATACCAGATAAG |
| N5A_Forward | CCATATGAATATTCCGGCGCAGATTACGGTTTTTAG |
| N5A_Reverse | CTAAAAACCGTAATCTGCGCCGGAATATTCATATGG |
| F58A_Forward | GCTTCCCTTAGCGATGCGGTTGATGGTTATTTAG |
| F58A_Reverse | CTAAATAACCATCAACCGCATCGCTAAGGGAAGC |
| R65A_Forward | GGTTATTTAGCTGCGAAATGGAATTTAG |
| R65A_Reverse | CTAAATTCCATTTCGCAGCTAAATAACC |
| K135A_Forward | GGTCAATTAGGTGCGATTAAAACAGC |
| K135A_Reverse | GCTGTTTTAATCGCACCTAATTGACC |
| V59D_Forward | CCCTTAGCGATTTTGATGATGGTTATTTAG |
| V59D_Reverse | CTAAATAACCATCATCAAAATCGCTAAGGG |
| V59N_Forward | CCCTTAGCGATTTTAATGATGGTTATTTAG |
| V59N_Reverse | CTAAATAACCATCATTAAAATCGCTAAGGG |
| G61S_Forward | CTTAGCGATTTTGTTGATAGCTATTTAGCTAGAAAATG |
| G61S_Reverse | CATTTTCTAGCTAAATAGCTATCAACAAAATCGCTAAG |
| A64V_Forward | GATGGTTATTTAGTGAGAAAATGGAA |
| A64V_Reverse | TTCCATTTTCTCACTAAATAACCATC |
| K75N_Forward | GTTACAAATATGGGGAATTTTTTGGATCCATTAG |
| K75N_Reverse | CTAATGGATCCAAAAAATTCCCCATATTTGTAAC |
| K135E_Forward | GTGCAGCTGGTCAATTAGGTGAAATTAAAACAGCAGTTAC |
| K135E_Reverse | GTAACTGCTGTTTTAATTTCACCTAATTGACCAGCTGCAC |

| | |
|---|---|
| S177F_Forward | GCGTTATTTTTACTATCTTATTTGGTATTGAATAC |
| S177F_Reverse | GTATTCAATACCAAATAAGATAGTAAAAATAACGC |
| D187E_Forward | CTTTTATAAAGGTAGAGAAGTTTTTAAACAAAAATAAC |
| D187E_Reverse | GTTATTTTTGTTTAAAAACTTCTCTACCTTTATAAAAG |
